# Normative models of enhancer function

**DOI:** 10.1101/2020.04.08.029405

**Authors:** Rok Grah, Benjamin Zoller, Gašper Tkačik

## Abstract

In prokaryotes, thermodynamic models of gene regulation provide a highly quantitative mapping from promoter sequences to gene expression levels that is compatible with *in vivo* and *in vitro* biophysical measurements. Such concordance has not been achieved for models of enhancer function in eukaryotes. In equilibrium models, it is difficult to reconcile the reported short transcription factor (TF) residence times on the DNA with the high specificity of regulation. In non-equilibrium models, progress is difficult due to an explosion in the number of parameters. Here, we navigate this complexity by looking for minimal non-equilibrium enhancer models that yield desired regulatory phenotypes: low TF residence time, high specificity and tunable cooperativity. We find that a single extra parameter, interpretable as the “linking rate” by which bound TFs interact with Mediator components, enables our models to escape equilibrium bounds and access optimal regulatory phenotypes, while remaining consistent with the reported phenomenology and simple enough to be inferred from upcoming experiments. We further find that high specificity in non-equilibrium models is in a tradeoff with gene expression noise, predicting bursty dynamics — an experimentally-observed hallmark of eukaryotic transcription. By drastically reducing the vast parameter space to a much smaller subspace that optimally realizes biological function prior to inference from data, our normative approach holds promise for mathematical models in systems biology.

---

An essential step in the control of eukaryotic gene expression is the interaction between transcription factors (TFs), various necessary co-factors, and TF binding sites (BSs) on the regulatory segments of DNA known as enhancers [1]. While we are far from having either a complete parts list for this extraordinarily complex regulatory machine or an insight into the dynamical interactions between its components, experimental observations have established a number of constraints on its operation: *(i)* TFs individually only recognize short, 6–10bp long binding site motifs [2]; *(ii)* TF residence times on the cognate binding sites can be as short as a few seconds and only 2–3 orders of magnitude longer than residence times on non-specific DNA [3–5]; *(iii)* the order of arrival of TFs to their binding sites can affect gene activation [4]; *(iv)* TFs do not activate transcription by RNA polymerase directly, but interact first with various co-activators, essential amongst which is the Mediator complex; *(v)* binding of multiple TFs is typically required within the same enhancer for its activation [6], which can lead to very precise downstream gene expression only in the presence of a specific combination of TF concentrations [7]; *(vi)* when activated, gene expression can be highly stochastic and bursty [8–10]; *(vii)* gene induction curves show varying degrees of steepness, suggesting tunable amounts of cooperativity among TFs [11]. Here we look for biophysical models of enhancer function consistent with these observations.

Mathematical modeling of gene regulation traces its origins to the paradigmatic examples of the *λ* bacteriophage switch [12] and the *lac* operon [13]. In prokaryotes, biophysical models have proven very successful [14–16], assuming gene expression to be proportional to the fraction of time RNA polymerase is bound to the promoter in thermodynamic equilibrium; TFs modulate this fraction via steric or energetic interactions with the polymerase. Crucially, these models are very compact: they are fully specified by enumerating all bound configurations and energies of the TFs and the polymerase on the promoter. While some open questions remain [17–19], the thermodynamic framework has provided a quantitative explanation for combinatorial regulation, cooperativity, and regulation by DNA looping [20, 21], while remaining consistent with experiments that also probe the kinetic rates [22, 23].

No such consensus framework exists for eukaryotic transcriptional control. Limited specificity of individual TFs *(i)* is hard to reconcile with the high specificity of regulation *(v)* and the suppression of regulatory crosstalk [24], suggesting non-equilibrium kinetic-proofreading schemes [25]. Likewise, short TF residence times *(ii)* and the importance of TF arrival ordering *(iii)* contradict the conceptual picture where stable enhanceosomes are assembled in equilibrium [4]. Kinetic schemes may be required to match the reported characteristics of bursty gene expression *(vi)* [26], or realize high cooperativity *(vii)* [27]. Thermodynamic models undisputedly have statistical power to predict expression from regulatory sequence even in eukaryotes [28], yet this does not resolve their biophysical inconsistencies or rule out non-equilibrium models. Unfortunately, mechanistically detailed non-equilibrium models entail an explosion in the complexity of the corresponding reaction schemes and the number of associated parameters: on the one hand, such models are intractable to infer from data, while on the other, it is difficult to understand which details are essential for the emergence of regulatory function.

To deal with this complexity, we systematically simplify the space of enhancer models. We adopt the normative approach, commonly encountered in the applications of optimality ideas in neuroscience and elsewhere [29–31]: we theoretically identify those models for which various performance measures of gene regulation, which we call “regulatory phenotypes”, are maximized. Such optimal model classes are our candidates that could subsequently be refined for particular biological systems and confronted with data. Thus, rather than inferring a single model from experimental data or constructing a complex, molecularly-detailed model for some specific enhancer, we find the simplest generalizations of the classic equilibrium regulatory schemes, such as Hill-type [32] or Monod-Wyman-Changeux regulation [33–35], to non-equilibrium processes, which drastically improves their regulatory performance while leaving the models simple to analyze, simulate, and fit to data.

## RESULTS

### A. Model

Multiple lines of evidence suggest that eukaryotic transcription is a two-state process which switches between active (ON) and inactive (OFF) states, with rates dependent on the transcription factor (TF) concentrations [36–38]. We sought to generalize classic regulatory schemes that can describe the balance between ON and OFF transcriptional states in equilibrium: a Hill-like scheme of “thermodynamic models” (discussed in SI Section 1.3), and a Monod-Wyman-Changeux-like (MWC) scheme introduced below.

Figure 1A shows a schematic of the proposed functional enhancer model (SI Section 1.1, see also Fig S1). A complex of transcriptional co-factors that we refer to as a “Mediator”^1^ can interact with TFs that bind and unbind from their DNA binding sites with baseline rates *k*_+_ and *k*_−_ (Fig 1B.i). Mediator – and thus the whole enhancer – can switch between its functional ON/OFF states with baseline rates *κ*_+_ and *κ*_−_ (Fig 1B.ii). Enhancer ON state and TF bound state are both stabilized (by a factor *α* relative to baseline rates) when a bound TF establishes a “link” with the Mediator (Fig 1B.iii). The molecular identity of such links can remain unspecified: it could, for example, correspond to an enzymatic creation of chemical marks (e.g., methylation, phosphorylation) on the TFs or Mediator proteins conditional on their physical proximity or interaction. Crucially, the links can be established and removed in processes that can break detailed balance and are thus out of equilibrium. Here, we consider that a link is established at a rate *k*_link_ between a bound TF and the Mediator complex; for simplicity, we assume that the links break when the TFs dissociate or upon the switch into OFF state (this assumption can be relaxed, see Fig S2).

**FIG. 1.**
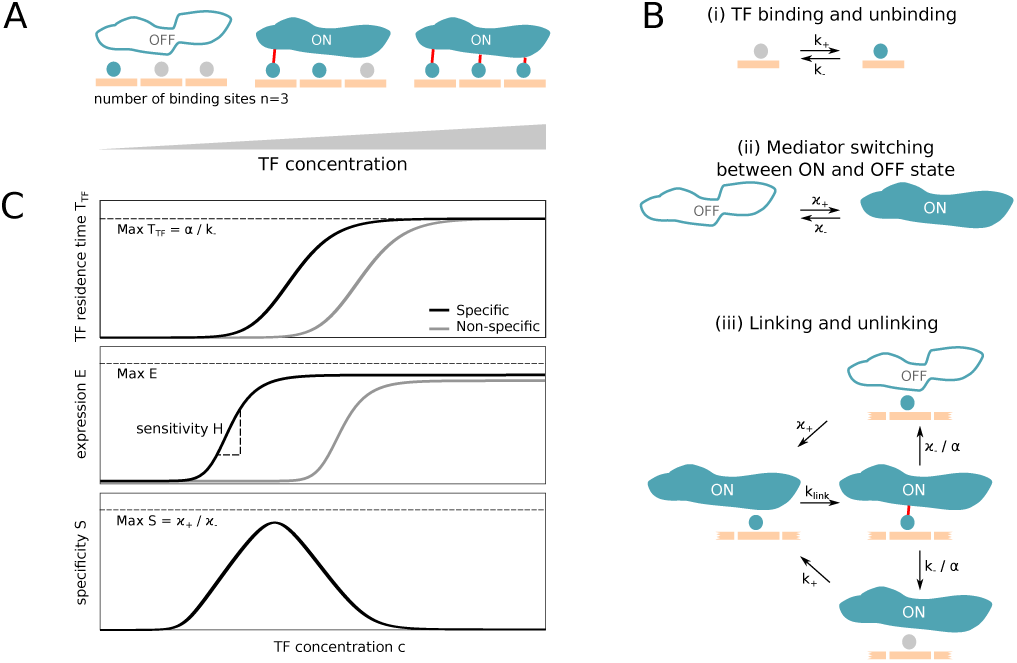
A non-equilibrium MWC-like model of enhancer function. **(A)** Schematic representation of transcription factors (TFs; tael circles) interacting with binding sites (BSs, here *n* = 3 orange slots) and the putative Mediator complex via links (red lines). The Mediator complex can be in two conformational states (OFF or ON), with the ON state enabling productive transcription of the regulated gene. Increasing TF concentration, *c*, facilitates TF binding and the switch into ON state (left-to-right). **(B)** Key reactions and rates of the non-equilibrium model. TFs can bind with concentration-dependent on rate 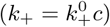 and unbind with basal rate *k*_−_ that is in principle sequence dependent (i). The Mediator state switches between the conformational states with basal rates *κ*_+_ and *κ*_−_ (ii). Linking and unlinking of TFs to Mediator (iii) can move the system out of equilibrium: links are established with rate *k*_link_, and the link stabilizes both TF residence and the ON state of the Mediator by a factor *α* per established link. **(C)** Regulatory phenotypes. Mean TF residence time, *T*_TF_, on specific sites in functional enhancers (black) vs random site on the DNA (gray) increases with concentration (top), as does mean expression, *E* (the fraction of time the Mediator is ON; induction curve, middle, with sensitivity, *H*, defined at mid-point expression). Specificity, *S*, is defined as the ratio of expression from the specific sites in the enhancer relative to the expression from random piece of DNA.

An important thrust of our investigations will concern the role of limited specificity of individual TFs to recognize their cognate sequences on the DNA. If sequence specificity arises primarily through TF binding −a strong, but relatively unchallenged assumption (that can also be relaxed within our framework, see Fig S3) − then we should ask how likely it is for the Mediator complex to form and activate at specific sites contained within functional enhancers (with low off-rates characteristic of strong eukaryotic TF binding sites, 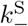) versus at random, non-specific sites on the DNA (with ∼2 orders-of-magnitude higher individual TF off-rates, 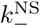) from which expression should not occur.

Given the number of TF binding sites (*n*) and the various rate parameters 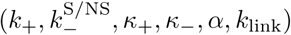 the full state of the system—i.e., the probability to observe any number of bound and/or linked TFs jointly with the ON/OFF state of the enhancer—evolves according to a Chemical Master Equation (SI Section 1.1) that can be solved exactly [39–41] or simulated using the Stochastic Simulation Algorithm [42]. Importantly, we show analytically that our scheme reduces to the true equilibrium MWC model in the limit *k*_link_ →∞: in this limit, there can be no distinction between a bound TF and a TF that is both bound and linked, and one can define a free energy *F* that governs the probability of enhancer being ON, which in our model is equal to (a normalized) mean expression level, *E* = *P*_ON_ = (1 + exp(*F*))^−1^, with

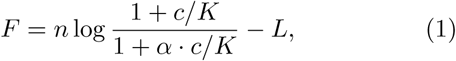

where 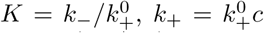 (see also Fig 1 caption), and *L* = log (*κ*_+_*/κ*_−_). The *k*_link_ parameter thus interpolates between the equilibrium limit in Eq (1), corresponding to a textbook MWC model, and various non-equilibrium (kinetic) schemes which we will explore next. A similar generalization with an equilibrium limit exists for thermodynamic Hill-type models, where, furthermore, *α* can be directly identified with cooperativity between DNA-bound TFs (see SI Section 1.3); we will see that this qualitative role of *α* will hold also for the MWC case.

### B. Regulatory phenotypes

How does the regulatory performance depend on the enhancer parameters and, in particular, on moving away from the equilibrium limit? To assess this question systematically, we define a number of “regulatory phenotypes”, enumerated in Table I and illustrated in Fig 1C. As a function of TF concentration, we compute: **(i)** individual TF residence time, *T*_TF_, on specific sites in functional enhancers, as well as on random, non-specific DNA, because these quantities have been experimentally reported in single-molecule experiments and provide strong constraints on enhancer function; **(ii)** average expression, *E*, for functional enhancers as well as random, non-specific DNA; we require *E* to be in the middle (∼0.5) of the wide range reported for functional enhancers; **(iii)** sensitivity of the induction curve at half-maximal induction, *H*, an observable quantity often interpreted as a signature of cooperativity in equilibrium models; **(iv)** specificity, *S*, as the ratio between expression *E* from functional enhancers vs from non-specific DNA, which should be as high as possible to prevent deleterious crosstalk or uncontrolled expression [24]; **(v)** expression noise, *N*, defined more precisely later, originating in stochastic enhancer ON/OFF switching.^2^

**TABLE I.** Regulatory phenotypes.

| Phenotype | Symbol | Value | Ref |
| --- | --- | --- | --- |
| TF residence time (specific BS) | $T_{\text{TF}}$ | $\sim 1 - 10$ s | [3, 43] |
| Expression (fraction of time ON) | $E$ | $0.01 - 0.9$ | [38, 44, 45] |
| Sensitivity (apparent Hill coef.) | $H$ | $1 - 10$ | [11] |
| Specificity | $S$ | — | — |
| Noise (std / mean protein exp.) | $N$ | $\sim 0.1 - 1$ | [46] |

### C. Specificity, residence time, and expression

Figure 2A explores the relationship between three regulatory phenotypes for a MWC-like enhancer scheme of Fig 1A: the average TF residence time (*T*_TF_), specificity (*S*), and the average expression (*E*), at fixed concentration *c*_0_ of the TFs. Each point in this “phase diagram” corresponds to a particular enhancer model; points are accessible by varying *α* and *k*_link_ (Fig 2B) and fall into a compact region that is bounded by intuitive, analytically-derivable limits to specificity and the residence time. As *α* tends to large values, *S* approaches 1, as it must: once a TF-Mediator complex forms, large *α* will ensure it never dissociates and expression *E* will tend to 1 (see also Fig 2D) irrespective of whether this occurred on a functional enhancer or a random piece of DNA – in this limit, all sequence discrimination ability is lost, yielding undesirable regulatory phenotypes. In contrast, the equilibrium (“EQ”) MWC limit as *k*_link_ → ∞ (Eq 1) is functional and, interestingly, corresponds to a non-monotonic curve in the phase diagram that lower-bounds the specificity of non-equilibrium (“NEQ”) models accessible at finite values of *k*_link_.

**FIG. 2.**
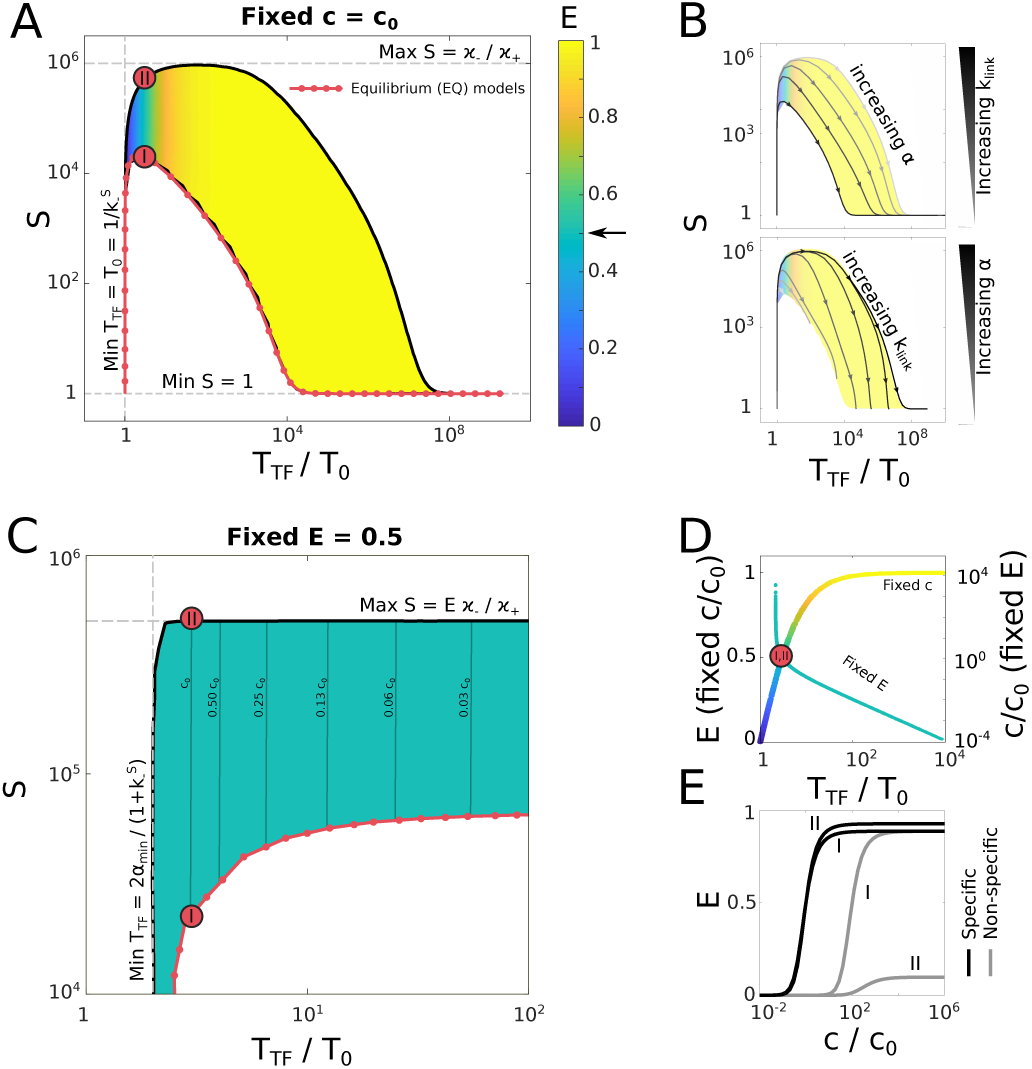
Accessible space of regulatory phenotypes. **(A)** Specificity, *S*, mean TF residence time, *T*_TF_ (expressed in units in inverse off-rate for isolated TFs at their specific sites, 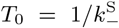, and average expression, *E* (color), for MWC-like models with *n* = 3 TF binding sites, obtained by varying *α* and *k*_link_ at fixed TF concentration, *c*_0_. Equilibrium models fall onto the red line; two models with equal TF residence times, I (EQ) and II (NEQ), are marked for comparison. Dashed gray lines show analytically-derived bounds. **(B)** Phase space of regulatory phenotypes is accessed by varying *α* at fixed values of *k*_link_ (grayscale; top) or varying *k*_link_ at fixed values of *α* (grayscale; bottom). **(C)** As in (A), but the TF concentration at each point in the phase space is adjusted to hold average expression fixed at *E* = 0.5 (green color). Plotted is a smaller region of phase space of interest; nearly vertical thin lines are equi-concentration contours (Fig S6). **(D)** All models in the phase diagrams in (A) and **(C)** approximately collapse onto nearly one-dimensional manifolds (“fixed c”, left axis, for (A); “fixed E”, right axis, for (C)) when plotted as a function of mean TF residence time, *T*_TF_, supporting the choice of this variable as a biologically-relevant observable. Color on the manifold corresponds to mean expression *E* using the colormap of (A). Vertical scales are chosen so that models I and II coincide. **(E)** Induction curves of equilibrium model I and non-equilibrium model II for expression from functional enhancer that contains specific sites (basal TF off-rate 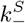; black curves) versus expression from random DNA containing non-specific sites (basal TF off-rate 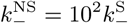 here; gray curves).

In a wide intermediate range of TF residence times, the full space of nonequilibrium MWC-like models—which we can exhaustively explore—offers large, orders-of-magnitude improvements in specificity, essentially utilizing a stochastic variant of Hopfield’s proofreading mechanism [25, 47]. This observation is generic, even though the precise values of *S* depend on parameters that we explore below, and *S* always remains bounded from above by *κ*_−_*/κ*_+_ (in equilibrium, this is related to stochastic, thermal-fluctuation-driven Mediator transitions to ON state even in absence of bound TFs). At the same average TF residence time and TF concentration, the best non-equilibrium model (II in Fig 2) will suppress expression from non-cognate DNA by almost two orders-of-magnitude relative to the best equilibrium model (I). These findings remain qualitatively unchanged for enhancers with larger number of binding sites (see Fig S4).

A comparison of various enhancer operating regimes is perhaps biologically more relevant at fixed mean expression, allowing the TF concentration to adjust accordingly under cells’ own control, as shown in Fig 2C for *E* = 0.5. As TF residence time lengthens with increasing *α*, TFs and the Mediator establish more stable complexes on the DNA and lower concentrations are needed for all models to reach the desired expression *E* (see also Fig 2D). Nevertheless, the ability of *α* to increase the specificity in equilibrium models is limited and saturates at a value substantially below the specificity reachable in nonequilibrium models at much smaller TF residence times. The observations of Figs 2A, C underscore an important, yet often overlooked, point: the ability to induce at low TF concentration (that is, high affinity) achieved through “cooperative interactions” at high *α* either has a detrimental, or, at best, a marginally beneficial effect for the ability to discriminate between cognate and random DNA sites (that is, high specificity) in equilibrium [24].

Figure 2E shows induction curves for expression from functional enhancers containing specific sites and from random DNA sites, for equilibrium (I) and non-equilibrium (II) models. Both yield essentially indistinguishable induction curves for expression from a functional enhancer (which is true generically across our phase diagram, see Fig S5), suggesting that it would be difficult to discriminate between the models based on induction curve measurements. In sharp contrast, the behavior of the two models is qualitatively different at non-specific DNA: with sufficiently high TF concentration (e.g., in an over-expression experiment), the EQ model I will fully induce even from random DNA as its binding sites get saturated by TFs; on the contrary, the nonequilibrium (NEQ) model II will start inducing at much higher *c*, and will never do so fully due to its proof-reading capability. Thus, given the relatively weak individual TF preference for cognate vs non-cognate DNA, one should look at the collective response of the gene expression machinery to mutated or random enhancer sequences for signatures of equilibrium vs non-equilibrium proofreading behavior.

### D. Sensitivity

Intuitively, sensitivity *H* measures the “steepness” of the induction curve. More precisely, *H* is proportional to the logartihmic derivative of the expression with log concentration at the point of half-maximal expression, so that for Hill-like functions, *E*(*c*) = *c*^*h*^*/*(*c*^*h*^ + *K*^*h*^), it corresponds exactly to the Hill coefficient, *H* = *h*. Figure 3A shows that *H* increases monotonically with *T*_TF_ (and thus with *α*, cf. Fig 2B), indicating that more stable TF-Mediator complexes indeed lead to higher apparent cooperativity, which is always upper-bounded by the number of TF binding sites in the enhancer, *n*. The highly-cooperative “enhanceosome” concept [48] would, in our framework, correspond to an equilibrium limit with very high *α*, and thus *H* ∼ *n*; yet the analysis above predicts vanishingly small specificity increases as this limit is approached. In contrast, we observe that the point at which the specificity advantage of nonequilibrium models is maximized, i.e., where *S*_NEQ_*/S*_EQ_ is largest, occurs far away from *H* = *n*, at much lower *H* values (Fig S8). If high specificity is biologically favored, we should therefore not expect the “number of known binding sites” to equal the “measured Hill coefficient of the induction curve” for well-functioning eukaryotic transcriptional schemes, even on theoretical grounds.

### E. Noise

Lastly, we turn our attention to gene expression noise. All stochastic two-state models have a steady state binomial variance of 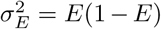 in enhancer state, where *E* is the probability of the enhancer to be ON. When ON, transcripts are made and subsequently translated into protein, which typically has a slow lifetime, *T*_*P*_, on the order of at least a few hours. Random fluctuations in enhancer state will cause random steady-state fluctuations in protein copy number around the average, *P*; these fluctuations can be quantified by noise, *N* = *σ*_*P*_ */P*. While there can be other contributions to noise (e.g., birth-death fluctuations due to protein production and degradation), we focus here solely on the effects of ON/OFF switching, since only these effects depend on the enhancer architecture [30].

How is noise in gene expression, *N*, related to the binomial variance, *σ*_*E*_? Based on simple noise propagation arguments [49, 50], fractional variance in protein should be equal to fractional variance in enhancer state times the noise filtering that depends on the timescales of enhancer switching, *T*_*E*_, and protein lifetime, *T*_*P*_ (here we assume *T*_*P*_ = 10 hours), so that *N* ^2^ = (*σ*_*P*_ */P*)^2^ ∼ (*σ*_*E*_*/E*)^2^ *T*_*E*_*/*(*T*_*E*_ + *T*_*P*_) (see SI Section 1.5 for exact derivation). Thus, if enhancer switches much faster than the protein lifetime, *T*_*E*_ « *T*_*P*_, protein dynamics almost entirely averages out the enhancer state fluctuations. Since all enhancer models have the same binomial variance, the gene expression noise in various models will be entirely determined by the mean expression, *E*, and the correlation time, *T*_*E*_, both of which we can compute analytically for any combination of enhancer model parameters in the phase diagram of Fig 2.

Figure 3B shows the phase diagram of accessible MWC-like regulatory phenotypes for the specificity (*S*), mean expression (*E*) and fraction of enhancer switching noise that propagates to gene expression, *T*_*E*_*/*(*T*_*E*_ + *T*_*P*_), found by varying *α* and *k*_link_. As in Fig 2, equilibrium models (“EQ”) have the lowest specificity *S*, but also lowest correlation time *T*_*E*_ and thus lowest noise, regardless of the average expression, *E*. There exist NEQ models that achieve higher specificity at a small increase in noise, but the highest specificity increases always come hand-in-hand with a substantial lengthening of the correlation times in enhancer state fluctuations, and thus with the inevitable increase in noise.

**FIG. 3.**
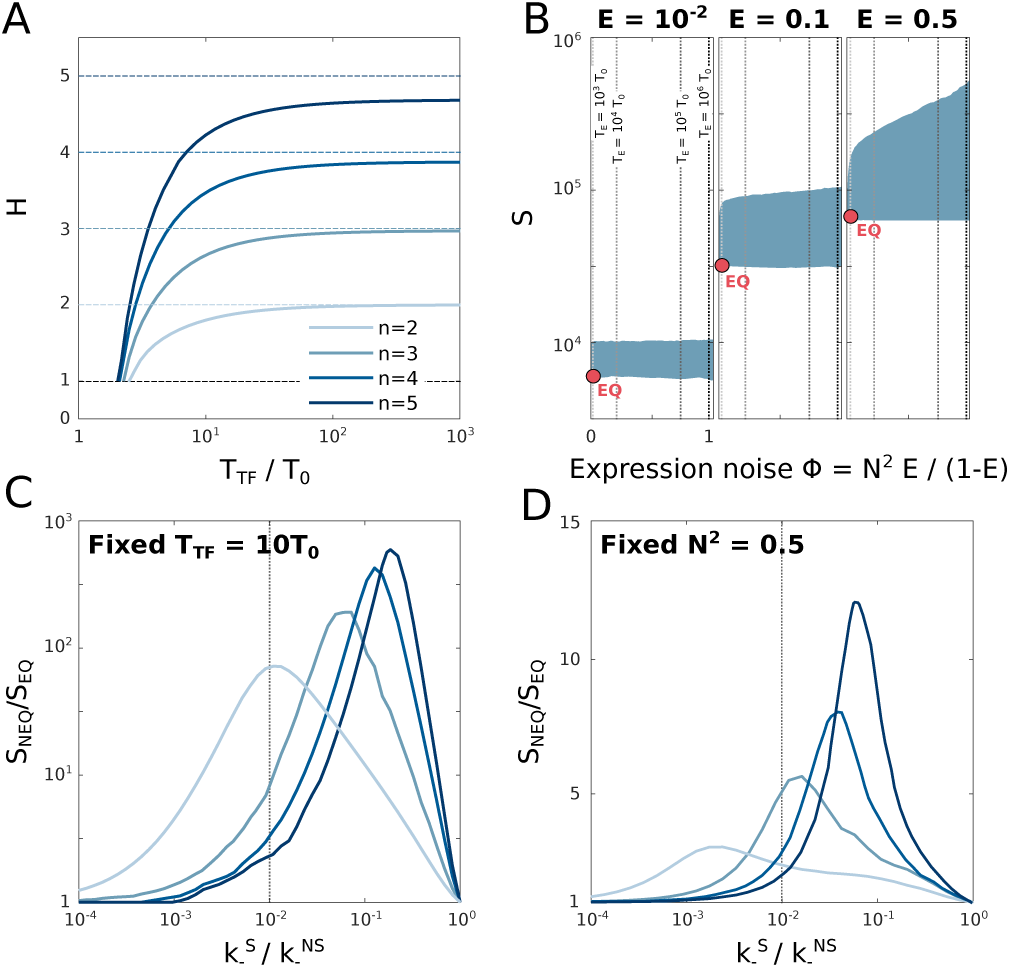
Limits to sensitivity and specificity. **(A)** Sensitivity (apparent Hill coefficient) *H* of enhancer models in the phase diagram of Fig 2C, at fixed mean expression, *E* = 0.5. All models collapse onto the manifolds shown for different number of TF binding sites, *n*. **(B)** Phase diagram of enhancer models for three different values of mean expression, *E* (columns), shows specificity *S* and fraction of variance in enhancer switching propagated to expression noise (see text). Compact blue region for each *E* shows all MWC-like models with *n* = 3 binding sites accessible by varying *α* and *k*_link_; equilibrium model (“EQ”) with lowest noise is shown as a red dot. Increase in noise is monotonically related to increase in enhancer correlation time, *T*_*E*_, marked with dashed vertical lines. Largest specificity increases over EQ models occur at high *T*_*E*_ and thus high noise (upper right corner of the blue region). **(C)** Maximal gain in enhancer specificity for non-equilibrium vs equilibrium models for different *n* (legend as in A), as a function of the intrinsic specificity of individual TF binding sites, 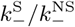. Expression is fixed to *E* = 0.5 and mean TF residence time to *T*_TF_*/T*_0_ = 10. Typical value 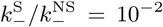 used in Fig 2 and panels A,B is shown in vertical dashed line. **(D)** Same as in (C), but with the comparison at fixed gene expression noise, *N* ^2^ = 0.5.

To better elucidate the tradeoffs and limits to specificity in non-equilibrium vs equilibrium models, we next explore how enhancer specificity gains depend on the ability of individual TFs to discriminate cognate binding sites from random DNA in Fig 3C. If individual TFs permit very strong discrimination 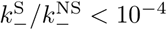; prokaryotic TF regime), NEQ models at fixed individual TF residence times, *T*_TF_, do not offer appreciable specificity increases in the collective enhancer response; in contrast, for the range around 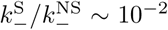 typically re-ported for eukaryotic TFs, the specificity increase ranges from ten to thousand-fold, with the peak depending on the number of TF binding sites, *n*, as well as baseline Mediator specificity limit, *κ*_−_*/κ*_+_ (as this increases, the peak specificity gain is higher and moves towards lower 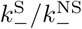, see Fig S9). If, instead of fixing 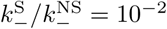 as we have done until now, we pick this ratio to maximize the specificity gain (*S*_NEQ_*/S*_EQ_) and again explore the noise-specificity tradeoff as in Fig 3B, we find that the extreme specificity gains are only possible when correlation times, *T*_*E*_ diverge (see Fig S10), implying high noise.

These observations are summarized in Fig 3D, showing the specificity gain of NEQ models relative to EQ models, if the comparison is made at fixed noise level rather than at fixed individual TF residence time as in Fig 3C. Specificity gains are limited to roughly ten-fold even when, as we do here, we systematically search for best NEQ models through the complete phase diagram in Fig 2C. The specificity-noise tradeoff thus appears unavoidable.

### F. Experimentally observable signatures of enhancer function

To illustrate how the proposed nonequilibrium (NEQ) MWC-like scheme could function in practice, we simulated it explicitly and compared it to an equilibrium (EQ) scheme with the same mean TF residence time in Fig 4. The two enhancers, composed of *n* = 5 TF binding sites, respond to a simulated protocol where the TF concentration is first switched from a minimal value that drives essentially no expression to a high value giving rise to *E* = 0.5, and after a long stationary period, the concentration is switched back to the low value. Figure 4A shows the occupancy of the binding sites and the functional ON/OFF state of the enhancer. Even though the two models share the same TF mean residence time and nearly indistinguishable induction curves (with *H* ∼ 2.7), their collective behaviors are markedly different: the EQ scheme appears to have significantly faster TF binding / unbinding as well as Mediator switching dynamics, whereas NEQ scheme undergoes long, “bursty” periods of sustained enhancer activation and TF binding that are punctuated by OFF periods. If the typical residence time of an isolated TF on its specific site were *T*_0_ = 1 s, NEQ enhancer could stay active even for hour-long periods (∼ 10^4^ s), just somewhat shorter than the protein lifetime (∼ 4 10^4^ s). Such enhancer-associated stable mediator clusters are consistent with recent experimental reports [51, 52].

**FIG. 4.**
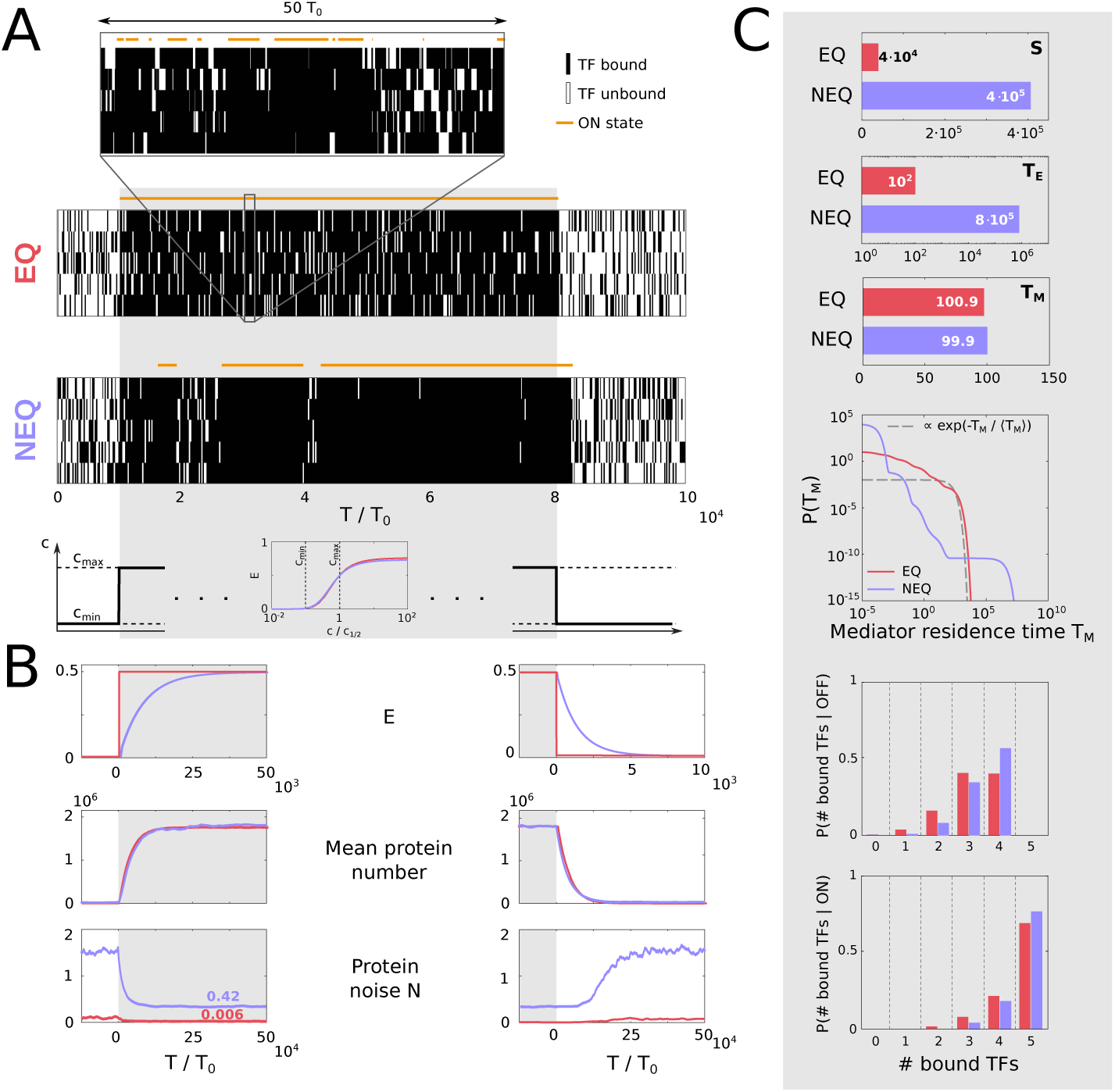
High-specificity non-equilibrium schemes predict bursty gene expression. **(A)** Stochastic simulation of an equilibrium (EQ) and a nonequilibrium (NEQ) enhancer model with *n* = 5 TF binding sites, responding to a TF concentration step (bottom-most panel). Average TF residence times are the matched between EQ and NEQ models at 2.1*T*_0_, 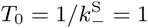s, and both induction curves (scaled for half-maximal concentration) are identical, with sensitivity *H* ≈ 2.7. When TF concentration is high, expression is fixed at *E* = 0.5. Parameters for NEQ model: *α* = 127, *k*_link_ = 2, *c*_max_ = 0.065; for EQ model: *k*_link_ *→ ∞, α* = 19.8,*c*_max_ = 0.037. Rasters show the occupancy of TF binding sites; orange line above shows the enhancer ON/OFF state; zoom-in for EQ model is necessary due to its fast dynamics. **(B)** Regulatory phenotypes for EQ and NEQ models during steady-state epoch (gray in A). Specificity (*S*) and enhancer state correlation time (*T*_*E*_) are higher for the NEQ model; the Mediator mean ON residence time, *T*_*M*_, is the same between the models, but the probability density function reveals a long tail in the NEQ scheme, and a nearly exponential distribution for the EQ scheme. Last two panels show the TF occupancy histogram during high TF concentration interval, conditional on the enhancer being OFF or ON. **(C)** Transient behavior of the mean enhancer state (*E*), mean protein number (*P*; assuming deterministic production/degradation protein dynamics given enhancer state), and gene expression noise, *N* = *σ*_*P*_ */P*, for the NEQ and EQ models, upon a TF concentration low-to-high switch (left column) and high-to-low switch (right column). Traces shown are computed as averages over 1000 stochastic simulation replicates.

The detailed steady-state behavior at high TF concentration is analyzed in Fig 4B. Consistent with our theoretical expectations, the NEQ scheme enables ten-fold higher specificity but at the cost of substantial noise in gene expression (*N* ∼ 0.42) due to strong transcriptional bursting. High noise is a direct consequence of the much longer correlation time of enhancer fluctuations, *T*_*E*_, for the NEQ scheme, seen in Fig 4A. Interestingly, the mean residence time of the enhancer ON state, *T*_*M*_, is nearly unchanged between the EQ and NEQ scheme at ∼ 100 s: but here, the mean turns to be a highly misleading statistic, as revealed by an in-depth exploration of the full probability density function. The NEQ scheme has a long tail of extended ON events interspersed with an excess of extremely short OFF events (due to high *κ*_−_ rate necessary for high specificity) relative to the EQ scheme (which, itself, does not deviate strongly from an exponential density function with a matched mean). The behavior of such an enhancer is highly cooperative even though the sensitivity (*H*) is not maximal: when the enhancer is ON, with very high probability all TFs are bound, and when OFF, often 4 out of 5 TFs are bound – yet the enhancer is not activated. In sum, a well-functioning non-equilibrium regulatory apparatus with its Mediator complex makes many short-lived attempts to switch ON, but only commits to a long, productive ON interval rarely and collectively, after insuring that activation is happening due to a sequence of valid molecular recognition events between several TFs and their cognate binding sites in a functional enhancer.

Transient behavior after a TF concentration change is analyzed in Fig 4C. The mean response time of the two models to the concentration change is governed by the correlation time of the enhancer state, *T*_*E*_, and is thus much slower for NEQ vs EQ models; but since the protein lifetime is even longer, the mean protein levels adjust equally quickly in the equilibrium and nonequilibrium cases. This suggests that the dynamics of the mean protein level is unlikely to discriminate between EQ and NEQ models. In contrast, live imaging of the nascent mRNA could put constraints on *T*_*E*_ [1]. In that case, the filtering time scale is the elongation time, typically on the order of a few minutes, while the reported transcriptional response times—and thus estimates of *T*_*E*_—would range from minutes to 1 − 2 hours [9, 26].

Steady-state noise levels at high induction, as reported already, are considerably higher for the NEQ model due to transcriptional bursting; an intriguing further suggestion of our analyses is a long transient in the noise levels upon a high-to-low TF concentration switch, which finally settles to a high fractional noise level (here, *N* ∼ 1.6) even at very low induction, due to sporadic transcriptional bursts.

## DISCUSSION

In this paper, we took a normative approach to address the complexity of eukaryotic gene regulatory schemes. We proposed a minimal extension to a well-known Monod-Wyman-Changeux model that can be applied to the switching between the active and inactive states of an enhancer. The one-parameter extension is kinetic and accesses nonequilibrium system behaviors. We analyzed the parameter space of the resulting model and visualized the phase diagram of “regulatory phenotypes”, quantities that are either experimentally constrained (such as mean expression, mean TF residence time, sensitivity), are likely to be optimized by evolutionary pressures (such as noise and specificity), or both. This allowed us to recognize and understand biophysical limits and trade-offs, and to identify the optimal operating regime of the proposed enhancer model that is consistent with current observations, as we summarize next.

Our analyses suggest the following: *(i)* individual TFs are limited in their ability to discriminate specific from random sites, 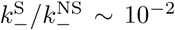, so high specificity must be a collective en−han−cer effect in the proofreading regime where 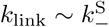; *(ii)* mean TF residence times in an enhancer are not m−uch higher than the typical TF residence time at an isolated specific site, *T*_*TF*_/*T*_0_ ≲ 10, enabling rapid turnover of bound TFs on the 1 − 10 s timescale; *(iii)* typical sensitivities are much lower than the total number of TF binding sites, yielding a reasonable specificity/noise balance at *H* ∼ *n/*2 (Fig S7,S8); *(iv)* Mediator basal rates should maximize *κ*_−_*/κ*_+_, i.e., mediator switches OFF essentially instantaneously if not stabilized by linked TFs; *(v)* TF concentrations required to activate the enhancer in this regime are substantially higher than expected for the equivalent but highly cooperative enhanceosome (at higher *α*); *(vi)* optimal nonequilibrium models achieve order-of-magnitude improvements in *S* relative to matched equilibrium models—thereby avoiding crosstalk and spurious gene expression—by suppressing induction from non-cognate (random) DNA, while induction curves from functional enhancers bear no clear signatures of non-equilibrium operation; *(vii)* to permit large increases in specificity *S*, enhancer state fluctuations will develop long timescale correlations, *T*_*E*_ » *T*_TF_ (but still be bounded by the protein lifetime, *T*_*E*_ ≲ *T*_*P*_ to enable noise averaging), leading to substantial observed noise levels; *(viii)* the enhancer ON residence time distribution will be non-exponential, with excess probability for very long-lived events, during which an enhancer could trigger a transcriptional burst following an interaction with the promoter; *(ix*) in our model, long correlation time, *T*_*E*_, in steady state also implies long (minutes to hours) response times when TF concentration change, which would be observable with live imaging on the transcriptional, but likely not protein-concentration, level.

We find it intriguing that a single-parameter extension of a classic equilibrium model led to such richness of observed behaviors, and to a suggestion that the optimal operating regime is very different from regulation at equilibrium. Central to this qualitative change is the fact that long fluctuation and response timescales of enhancer activation appear necessary to achieve high specificity of regulation through proofreading. Such long timescales are not inconsistent with our current knowledge. Indeed, some developmental enhancers form active clusters (super-enhancers) that are rather long-lived (order of minute to hours), perhaps precisely because developmental events need to be guided with extraordinary precision [52, 53].

A strong objection to our model could be that it is too simple: after all, we neglected many structural and molecular details, many of which we may not even know yet. This is certainly true and was done, in part, on purpose, to permit exhaustive analysis across the complete parameter space. Such understanding would have been impossible if we explored much richer models or were concerned with quantitative fitting to a particular dataset. These are clearly the next steps, to which we contribute by highlighting the functional importance of breaking the equilibrium link between TF binding and enhancer activation state. Since our model is fully probabilistic, specializing it for a particular experimental setup, e.g., live transcriptional imaging, and doing rigorous inference is technically tractable, but beyond the scope of this paper.

Perhaps a key simplification of our model is the link between enhancer / Mediator ON state and transcriptional activity. We assumed that expression is proportional to the probability of enhancer state to be ON, yet the enhancer-promoter interaction itself is a matter of vibrant current experimentation and modeling [10, 51, 54–56]. For example, long-lived activated enhancers that we predict could interact with promoters only intermittently to trigger transcriptional bursts, as suggested by the “dynamic kissing model” [52], which could substantially impact the experimentally-observable quantitative noise signatures of enhancer function at the transcriptional level. Whatever the true nature of enhancer-promoter interactions might be, however, they are unlikely to be able to remove excess enhancer switching noise, due to its slow timescale, suggesting that the tradeoffs that we identify should hold generically.

One could also question whether the importance we ascribed to high specificity is really warranted. Evolutionarily, regulatory crosstalk due to lower specificity helps networks evolve during transient bouts of adaptation, even though it could be ultimately selected against [57]. Mechanistically, molecular mechanisms such as chromatin modification or the regulated 3D structure of DNA decrease the number of possible non-cognate targets that could trigger erroneous gene expression [58, 59], and thus alleviate the need for the high specificity of the transcriptional control. Empirically, there is ample evidence for abortive or non-sensical transcriptional activity [60, 61], whose products could be dealt with downstream or simply ignored by the cell. Yet it is also clear that regulatory specificity must be a collective effect, as individual TFs bind pervasively across DNA even in non-regulatory regions [62], and self-consistent arguments suggest that in absence of non-equilibrium mechanisms, crosstalk could be overwhelming in eukaryotes [24]. It is also possible that real enhancers are very diverse with large variation along the specificity axis, thereby navigating the noise-specificity tradeoff as appropriate given the biological context. Where some erroneous induction can be tolerated, expression could be quicker, less noisy, and closer to equilibrium. In contrast, where tight control is needed, enhancers could take a substantial amount of time to commit to expression correctly, perhaps benefitting additionally from extra time-averaging that could further reduce the Berg-Purcell-type noise intrinsic to TF concentration sensing [50, 63–65].

## ACKNOWLEDGMENTS

GT acknowledges the support of the Human Frontiers Science Program RGP0034/2018. RG was supported by the Austrian Academy of Sciences DOC fellowship. RG thanks S. Avvakumov for helpful discussions.

## Footnotes

1 Our nomenclature is simply a shorthand for all co-factors necessary for eukaryotic transcriptional activation at an enhancer, which can include proteins not strictly a part of the Mediator family.

2 Protein noise levels in Table I are estimated from reported mRNA noise levels.

