## Supporting Information for "Normative models of enhancer function"

R Grah, B Zoller, G Tkačik

**2 SI Figures** **28**

### 1 Model

#### 1.1 Model specification

We consider a class of toy models for transcriptional regulation that could plausibly be employed by eukaryotic cells. Specifically, we look for models where transcription factors (TFs) interact with special regulatory sequences on the DNA, known as binding sites (BSs) in the enhancer, to control the expression of a given gene. The emphasis here is not on devising a single scheme that has a direct molecular interpretation, but rather to ask about *possible* schemes that simultaneously achieve several properties which are desirable for efficient regulation and are consistent with metazoan observations.

Our model, schematically displayed in Fig S1 for a single binding site for simplicity, includes a large pool of TFs at a fixed concentration $c$ of the same type that can bind to $n$ binding sites in the enhancer with a concentration-dependent rate $k_+ = k_+^0 c$ and unbind with a concentration-independent rate $k_-$. Additionally, a complex of transcriptional cofactors that we refer to as a "Mediator" can switch between `ON` and `OFF` state with rates $\kappa_+$ and $\kappa_-$. Only when Mediator is found in an `ON` state, can a so-called link between any bound TF and Mediator be established with a rate $k_{\text{link}}$. In principle, the link could be removed actively at a rate $k_{\text{unlink}}$, but here we assume for simplicity that $k_{\text{unlink}} = 0$ (we later relax this assumption). While the links are not removed actively, they are removed automatically when the TFs dissociate or upon the Mediator switch into `OFF` state. Molecularly, the link formation and removal could be catalyzed by dedicated enzymes, and when coupled to an energy source, could be kept out-of-equilibrium. The formation of any link increases the stability of the linked complex by decreasing the rate of unbinding of the linked molecules (and the rate of Mediator switching `OFF`) by a constant multiplicative factor; in other words, the Mediator `OFF` switching rate falls as a power in the number of links with TFs. We will see that this ansatz permits our model to have a clear thermodynamic-equilibrium limit.

**Parameters of the model.** The parameters of the model are:

- $n$ – number of specific binding sites in the enhancer.
- $\alpha$ – fold-reduction in the unbinding rate of a linked TF ($k_- \to k_-/\alpha$) or Mediator `OFF`

switching rate ($\kappa_- \to \kappa_-/\alpha^b$, $b \in \{0, 1, \ldots, n\}$ is number of links Mediator has with bound TFs). We focus on values $\alpha \geqslant 1$ which stabilize the bound complexes.

- $k_+$ – binding rate of TF to a BS. $k_+ = k_+^0 c$, with $k_+^0$ being the binding rate per unit concentration and $c$ is the free concentration of TFs.

- $k_-$ – unbinding rate of TF bound in the enhancer. Generally, the unbinding rate of a TF depends on the presence of a link of the TF with the Mediator. If a link is present, the unbinding rate becomes $k_-/\alpha$. Importantly, $k_-$ is the only sequence dependent quantity in the model: $k_- = k_-^{\mathrm{S}}$ for an isolated unlinked TF bound on the specific BS, and $k_- = k_-^{\mathrm{NS}}$ for an isolated unlinked TF bound on a non-specific, random site.

- $k_{\mathrm{link}}$ – rate of establishing new links between bound TF and the Mediator in `ON` state. If the link-forming reaction were catalyzed by an enzyme with own sequence specificity, we can achieve an even higher specificity of our regulatory scheme (examined later in this document). To be conservative, we set this rate to be constant and thus have no sequence specificity, i.e., once TF is bound and Mediator is in `ON` state, the link can be created with the same rate regardless of whether this happens in the enhancer or on a random piece of genomic sequence.

- $\kappa_+$ – switching rate into `ON` state of the Mediator.

- $\kappa_-$ – switching rate into `OFF` state of the Mediator. Generally, this rate depends on the number of links a Mediator has established with TFs. Thus, this rate takes a form $\kappa_-/\alpha^b$, where $b$ is the number of linked bound TFs.

- $k_{\mathrm{unlink}}$ – rate of active link removal. In our default model, we set this parameter to $k_{\mathrm{unlink}} = 0$. Fig S2 analyzes the effects of this assumption.

In our analysis we will focus on the effects of $k_{\mathrm{link}}$, $\alpha$, and $n$ while taking representative values of the other parameters; if not stated otherwise, we use $k_{\mathrm{unlink}} = 0$, $k_+ = ck_+^0$, with $c = c_0 = 0.01$ and $k_+^0 = 1$, $\kappa_- = 10^4$, $\kappa_+ = 10^{-2}$, $k_-^{\mathrm{S}} = 10^{-2}$, and $k_-^{\mathrm{NS}} = 1$. The latter also sets the timescale in our model, that is, we define $T_0 = 1/k_-^{\mathrm{NS}} = 1$, i.e., the typical time a TF in isolation is bound on a random, non-specific site on the DNA, as our time unit. For exploration of the phase space we use $\alpha \in (1, 10^{10})$ and $k_{\mathrm{link}} \in (10^{-8}, 10^8)$.

**Dynamical variables and computation of the model.** The dynamical variables of our model are:

- $s_i$ – an indicator variable in $\{0, 1\}$ indicating if a TF is bound on the site $i = 1, \ldots, n$.
- $b_i$ – an indicator variable in $\{0, 1\}$ indicating if TF at site $i$ has a link with the Mediator. It can take a value of 1 only if the TF at site $i$ is bound, i.e., if $s_i = 1$.
- $s_M$ – an indicator variable in $\{0, 1\}$ indicating if the Mediator is in `ON` state.

The behavior of the system in state space of $\{s_i, b_i, s_M\}$ is a continuous-time Markov chain, with the rates fixed by our parameters. Generally, we can write down our system as a Master equation for the Markov chain:

$$\frac{\mathrm{d}\mathbf{V}}{\mathrm{d}t} = \hat{M}\mathbf{V}, \tag{S1}$$

with $\mathbf{V}$ being a vector whose components are the probabilities of the system to be in any of the states at time $t$ (and thus $\sum_{j=1}^{m} V_j = 1$, where the sum is taken over all $m$ components of the vector $\mathbf{V}$) and $\hat{M}$ the transition matrix between different states. However, in practice the number of all possible microstates is large, making explicit manipulation of the Master equation feasible only for smaller values of $n$.

**Constructing the transition matrix $\hat{M}$.** In this paragraph we describe how to construct the transition matrix $\hat{M}$. First, we define a state vector $\mathbf{B} = (s_M, s_1, \ldots, s_n, b_1, \ldots, b_n)$. With the state vector $\mathbf{B}$ we can enumerate all possible states of $s_M \in \{0, 1\}$, $s_i \in \{0, 1\}$, and $b_i \in \{0, 1\}$. However, the $b_i$ values are constrained by the binding state of the TFs ($s_i$) and cannot independently take on arbitrary values. For example, if $s_i = 0$ ($i$–th TF not bound), then there can never be any link, i.e., $b_i = 0$. Only if $s_i = 1$, then $b_i \in \{0, 1\}$. If we take the three example from Fig 1A with $n = 3$ at increasing TF concentrations, the corresponding state vectors would be: $S_{\text{low c}} = (0, 1, 0, 0, 0, 0, 0)$, $S_{\text{medium c}} = (1, 1, 1, 0, 1, 0, 0)$, $S_{\text{high c}} = (1, 1, 1, 1, 1, 1, 1)$.

Altogether there are

$$m = \sum_{i=0}^{n} \binom{n}{i} 2^i + 2^n \tag{S2}$$

different states, where the sum goes over all possible combinations of $i$ bound and potentially linked TFs with the Mediator in `ON` state. The second part represents the number of different states when Mediator is in `OFF` state, i.e., when $s_M = 0$. Next, we order the states such that states with $s_M = 1$ come first, followed by states $s_M = 0$.

We can write the transition matrix by accounting for all possible events, and then finding states between which events cannot occur. Roughly, there are 3 types of events: (i) binding and unbinding of TFs, (ii) linking and unlinking, and (iii) switching the Mediator between `ON` and `OFF` (main text Fig 1B). In the following procedure, we will go over the three different types of events, finding all possible transitions between them, and assigning rates to those state-change events in the transition matrix. Due to symmetries, directly finding only a subset of events is enough. For example, starting at state $j$, let us assume that a linking event can lead to state $k$. As the unlinking events are reciprocal to linking events, the unlinking of the same TF (assuming states of all other TFs and Mediator did not change) would lead from state $k$ to state $j$. However, this is not entirely correct for binding and unbinding events $-$ the extra complication is that unbinding can also destroy a link and reciprocity between binding and unbinding does not always exist. Therefore, if unbinding of a TF leads from state $j$ to state $k$, the state $j$ can be reached from state $k$ only if the unbinding did not destroy a link. This means that there are more unbinding transitions than there are binding transitions. Using this approach, we will find all possible state transitions. The procedure to write down the elements in the transition matrix $\hat{M}$ is:

For each possible state vector **B** (`original state`) do

1. First we locate all linking and unlinking transitions. Therefore, for each TF $i$ in order:
   - If the $i-$th TF is not linked (i.e., $B(i + n + 1) = 0$), continue to the next TF.
   - Otherwise, define a `new state` by removing the present link at $i-$th TF from the

original state. This new state has all elements the same as original state with the exception of no link at TF $i$ ($B_{\text{new}}(i + n + 1) = 0$).

- Mark the unlinking transition with unlinking rate as $M(\texttt{new state}, \texttt{original state}) = k_{\text{unlink}}$.
- Mark linking transition as $M(\texttt{original state}, \texttt{new state}) = k_{\text{link}}$.

2. Next, we locate all binding and unbinding transitions. As the two are not always reciprocal, we have to follow if a link is broken when unbinding occurs. For each TF $i$ in order:

   - If the $i-$th TF is not bound ($B(i + 1) = 0$), continue to the next TF.
   - Otherwise, define a `new state` by removing the bound $i-$th TF from the `original state`; further, remove the link of $i-$th TF (if it existed in the `original state`).
   - Count the number of removed links $b_i$: 0 if the bound TF was not linked and 1 if it was.
   - Mark the unbinding transition with unbinding rate as $M(\texttt{new state}, \texttt{original state}) = k_-/\alpha_i^b$.
   - If no link was removed (i.e., $b_i = 0$), this means that binding from the new to the original state can occur. Therefore, mark binding transition as $M(\texttt{original state}, \texttt{new state}) = k_+$.

3. Lastly, locate the states that are affected by Mediator switching:

   - If the Mediator is `OFF` in the `original state`, continue to the next state vector $\mathbf{B}$ and restart this processing at (1). If the Mediator is `ON` in the `original state`, define a `new state` by switching the Mediator into `OFF` state.
   - If any links existed between the Mediator and any other TF in the `original state`, remove them in the `new state`.
   - Count the number of removed links $b$.
   - Mark the transition into `OFF` state as: $M(\texttt{new state}, \texttt{original state}) = \kappa_-/\alpha^b$.
   - If no links were removed (i.e., $b = 0$), mark the transition into `ON` state as: $M(\texttt{original state}, \texttt{new state}) = \kappa_+$.

At the end we set the diagonal values as minus sum of the columns.

#### 1.2 Residence time distributions

Since all the individual processes involved in our enhancer models are Poisson processes occurring either sequentially or in phase, the residence time distributions of a given TF site or Mediator being `ON` are phase-type distributions. To compute these distributions, we first need to define various subsets of states. The set $I_b$ of states correspond to a given TF site or Mediator being bound (`ON`). The set $I_u$ of states correspond to a given TF site or Mediator being unbound (`OFF`). The set $I_{bn}$ of states correspond to a given TF site or Mediator being bound (`ON`) with no link attached. We can then define the following matrix from the original transition matrix $\hat{M}$ (Eq. S1)

$$\hat{M}_w = \hat{J}\hat{M}\hat{J}^{\mathbf{t}}, \tag{S3}$$

where $\hat{J}$ is a diagonal matrix whose entries $J_{ii}$ are equal to 1 if $i \in I_b$ and zero otherwise. We will also define a vector $\mathbf{a}$ describing the probability for the system to have just settled in any of the bound states

$$\mathbf{a}_i = \frac{\tilde{\mathbf{a}}_i}{\sum_{i=1}^{n} \tilde{\mathbf{a}}_i} \quad \text{with} \quad \tilde{\mathbf{a}}_i = \begin{cases} \sum_{j \in I_u} \hat{M}_{ij}\mathbf{V}_j & \text{for } i \in I_{bn} \\ 0 & \text{otherwise} \end{cases}, \tag{S4}$$

where $\mathbf{V}$ is the vector of steady state occupancies that is computed from Eq. S31. The probability density function for the residence time of a given TF site or Mediator being bound is then given by

$$f(T) = -\mathbf{I}^{\mathbf{t}} \exp(\hat{M}_w T)\hat{M}_w \mathbf{a}, \tag{S5}$$

where $\mathbf{I}$ is a vector whose entries are all equal to one. Of note, the exponential here is the matrix exponential. The mean residence time $\mu_T$ and the variance of the distribution $\sigma_T^2$ are then given by

$$\begin{aligned} \mu_T &= -\mathbf{I}^{\mathbf{t}} \hat{M}_w^{-1}\mathbf{a} \qquad \text{(S6)} \\ \sigma_T^2 &= 2\mathbf{I}^{\mathbf{t}} \hat{M}_w^{-2}\mathbf{a} - \mu_T^2. \end{aligned}$$

#### 1.3 Equilibrium limits

Our model is a generalization of an equilibrium MWC model. In the following section, we first show that our model reduces to the MWC model in the equilibrium limit, and we then derive the different regulatory phenotypes in this limit. At the end we also address the Hill-type models.

**MWC model.** As a thermodynamic equilibrium model, one can fully specify the MWC model by means of a partition function $Z$ that enumerates all the possible states of the system. The partition function of the MWC model with $n$ TF binding sites can be written as

$$Z = \sum_{\{\sigma_M, \sigma_i\}} \exp\left[L\sigma_M + (\epsilon + \log c + \delta\sigma_M)\sum_{i=1}^{n}\sigma_i\right], \tag{S7}$$

where $\sigma_M \in \{0,1\}$ and $\sigma_i \in \{0,1\}$ are the occupancy variables for Mediator and the TF binding sites $i$. The different energy contributions in our model are $L$, $\epsilon$ and $\delta$, which represent the energy difference between the Mediator `ON` and `OFF` state, between an empty and occupied TF binding site, and the energy benefit due to the established link. Their Boltzmann states can be respectively written as $e^L$, $e^\delta$, and $ce^\epsilon$ where $c$ represents the concentration of TFs. For example, the energy and Bolztmann weight of a state with Mediator in `ON` state and two bound and linked TFs would read $L + 2\epsilon + 2\delta$ and $ce^{L+2\epsilon+2\delta}$, respectively.

**MWC is an equilibrium limit of our model.** It turns out that our proposed model collapses into an equilibrium MWC model when taking the limit $k_{\text{link}} \to \infty$. This can be shown without loss of generality, by examining the simplest case of our model with $n = 1$. In that case, the single irreversible step dictating the non-equilibrium nature of our model occurs between the two following states: i) the TF is bound and Mediator `ON` but no link is present, and ii) a link is established between Mediator and the TF (Fig S.I). When increasing $k_{\text{link}}$, such that $k_{\text{link}} \gg k_-, \kappa_-$, the transition between these two states becomes very fast, and the dwell time in the first state becomes negligible. Thus, in the limit of $k_{\text{link}} \to \infty$, the two states with a bound TF and Mediator `ON` collapse into a single state where a link is always present (Fig S.I). The resulting kinetic scheme exactly corresponds to an equilibrium MWC model.

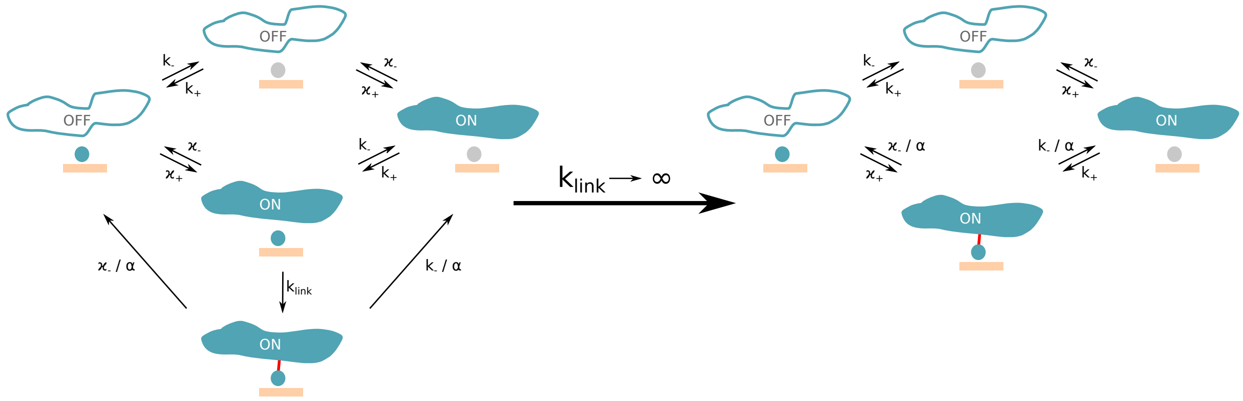

Fig S.I: **Our model is a generalization of MWC model.** A comparison between schemes for our non-equilibrium model (left) and MWC model (right) for $n = 1$. When taking the limit $k_{\text{link}} \to \infty$, our model converges into MWC model.

To make the correspondence clear, we connect the equilibrium energies defined in the MWC partition function (Eq. S7) with the kinetic rates of our model. Due to equilibrium, all processes follow detailed balance:

$$\pi_i W_{ij} = \pi_j W_{ji}, \tag{S8}$$

where $W_{ij}$ is the transition rate from state $i$ to state $j$, and $\pi_i$ and $\pi_j$ are the equilibrium probabilities of being in states $i$ and $j$, respectively.

First, we compare two states devoid of bound TFs where Mediator is `OFF` and `ON` respectively. Transitioning between these two states only involves the Mediator kinetic rates $\kappa_+$ and $\kappa_-$. Applying detailed balance gives $\frac{1}{Z}\kappa_+ = \frac{e^L}{Z}\kappa_-$, where $1/Z$ and $e^L/Z$ are the equilibrium probabilities $\pi$ of the `ON` and `OFF` state, respectively. It follows from this condition that $e^L = \frac{\kappa_+}{\kappa_-}$.

Similarly, we compare transitions between an empty state and a state with one bound TF. Following detailed balance we obtain $\frac{1}{Z}k_+ = \frac{ce^\epsilon}{Z}k_-$, where $1/Z$ and $\frac{ce^\epsilon}{Z}$ are the equilibrium probabilities $\pi$ of the TF unbound and bound state. The transition rates $k_+$ and $k_-$ are the rates of TF binding and unbinding, respectively. It thus follows that $ce^\epsilon = \frac{k_+}{k_-}$.

Lastly, we compare the two following states; i) Mediator is ON without any TF bound, and ii) Mediator is ON and a TF is bound. In the equilibrium limit, both Mediator and the TF are linked when present together. The equilibrium probabilities $\pi$ of the two states are $ce^L/Z$ and $ce^{L+\epsilon+\delta}/Z$, respectively. The transition rates between these states are $k_+$ and $k_-/\alpha$. By applying the detailed balance condition, $\frac{ce^L}{Z}k_+ = \frac{ce^{L+\epsilon+\delta}}{Z}\frac{k_-}{\alpha}$, we obtain $e^\delta = \alpha$. This demonstrates a one-to-one correspondence between the "cooperative energy of binding" in thermodynamic models of gene regulation, and the parameter $\alpha$ of the non-equilibrium model.

**Expression.** In our model, we defined expression as the occupancy of Mediator in the ON state. To calculate the expected expression in the MWC model, we first separate the patition function (Eq. S7) in two sub-partitions $Z_{\text{ON}}$ and $Z_{\text{OFF}}$ such that $Z = Z_{\text{ON}} + Z_{\text{OFF}}$. Here, $Z_{\text{ON}}$ and $Z_{\text{OFF}}$ correspond to the sum over all the states with Mediator ON and OFF respectively. These sums can be calculated as follows:

$$\begin{aligned} Z_{\text{ON}} =& e^L \sum_{\{\sigma_i\}} \exp\left[(\epsilon + \log c + \delta)\sum_{i=1}^{n}\sigma_i\right] \\ =& e^L \sum_{\{\sigma_i\}} \prod_{i=1}^{n} ce^\epsilon e^\delta e^{\sigma_i} \\ =& e^L \sum_{k=0}^{n} \binom{n}{k} (ce^\epsilon e^\delta)^k \cdot 1^{n-k} \\ =& e^L (1 + ce^\epsilon e^\delta)^n. \end{aligned} \tag{S9}$$

Similarly, we obtain $Z_{\text{OFF}} = (1 + ce^\epsilon)^n$. The probabilities to find the Mediator in the ON and OFF states can then be expressed as

$$\begin{aligned} P_{\text{ON}} &= \frac{Z_{\text{ON}}}{Z} = \frac{1}{Z} e^L \left(1 + ce^\epsilon e^\delta\right)^n \\ P_{\text{OFF}} &= \frac{Z_{\text{OFF}}}{Z} = \frac{1}{Z} (1 + ce^\epsilon)^n . \end{aligned} \tag{S10}$$

We can thus write the occupancy of Mediator in the `ON` state, which corresponds to our definition of expression:

$$E = \frac{P_{\mathrm{ON}}}{P_{\mathrm{ON}} + P_{\mathrm{OFF}}} = \frac{e^L \left(1 + ce^{\epsilon}e^{\delta}\right)^n}{e^L \left(1 + ce^{\epsilon}e^{\delta}\right)^n + \left(1 + ce^{\epsilon}\right)^n} = \left[1 + e^{-L}\left(\frac{1 + ce^{\epsilon}}{1 + ce^{\epsilon}e^{\delta}}\right)^n\right]^{-1}. \tag{S11}$$

As in the main text, all the bounds derived and reported below come from varying $k_{\mathrm{link}}$ and $\alpha$, while keeping other parameters constant. The only exception is concentration $c$ which is either kept constant or adjusted to achieve fixed expression $E$.

As per definition of occupancy, expression is bounded from above by $E = 1$; that occurs when Mediator is always in `ON` state. The lower bound, $\min E$, occurs when binding sites are almost never occupied. In that case, the expression is solely determined by the intrinsic Mediator `ON` probability, thus $E = \kappa_+/(\kappa_+ + \kappa_-)$, which reduces to $E = \kappa_+/\kappa_-$ when $\kappa_- \gg \kappa_+$, as we assumed. Therefore, expression is limited to $E \in (\frac{\kappa_+}{\kappa_-}, 1)$.

When fixing specific expression to $E^{\mathrm{S}} = E_0$, concentration must vary to meet that requirement. By equating $E_0 = \frac{P_{\mathrm{ON}}}{P_{\mathrm{ON}} + P_{\mathrm{OFF}}}$, and solving for $c$, we obtain:

$$c = \frac{k_-}{k_+^0}\frac{x - 1}{1 - x\alpha} \quad \text{with} \quad x = \left(\frac{\kappa_+(1 - E_0)}{\kappa_- E_0}\right)^{1/n}. \tag{S12}$$

Since concentration must be positive, $\alpha$ must satisfy $\alpha \geqslant \frac{1}{x}$. From this inequality we obtain a lower bound for $\alpha$

$$\alpha_{\min} = \left(\frac{\kappa_- E_0}{\kappa_+(1 - E_0)}\right)^{1/n}. \tag{S13}$$

.

**Specificity.** We define specificity as the ratio of expression from a functional enhancer, $E^{\mathrm{S}}$, and expression from a random piece of sequence, $E^{\mathrm{NS}}$. Therefore:

$$S = \frac{E^{\mathrm{S}}}{E^{\mathrm{NS}}}. \tag{S14}$$

By definition of specific binding site, $E^{\mathrm{S}} \geqslant E^{\mathrm{NS}}$, leading to $\min S = 1$.

Furthermore, a general upper bound of specificity is given by the ratio of Mediator switching rates, $\max S = \kappa_-/\kappa_+$. This upper bound is attained when specific expression is maximal, $E^{\mathrm{S}} = 1$, while non-specific expression is minimal $E^{\mathrm{NS}} = \frac{\kappa_+}{\kappa_-}$. Thus, $S \in (1, \kappa_-/\kappa_+)$. However, if a system has a fixed specific expression $E \neq 1$ (as in Fig. 2C), the upper bound is adjusted by a factor of $E$. Indeed, the specific expression takes value $E^{\mathrm{S}} = E$ by construction while minimal non-specific expression is again $E^{\mathrm{NS}} = \kappa_+/\kappa_-$. Taking their ratio we thus obtain $\max S_{\text{fixed E}} = E\kappa_-/\kappa_+$.

**Residence time.** We defined TF residence time as the average time a TF spends bound to its specific binding site. To calculate the TF residence time, we assumed that changes in the Mediator state do not happen while the TF is bound, which means that the residence time of TFs is either much shorter or longer than the time Mediator spends in `ON` state. This is a valid assumption in our model, since the Mediator `ON` state is either very short lived due to high `OFF` rate in absence of any link, or very long lived due to very small `OFF` rate in presence of stabilizing links.

Based on the assumption above, we can calculate the residence time as the weighted average of the average time spent in the two following configurations: a TF resides on a binding site with a link (Mediator `ON`), and without any link (Mediator `OFF`). These average times are given by the inverse of the escape rate, namely the inverse TF unbinding rate $e^{\delta}/k_-$ with a link and $1/k_-$ without a link. To obtain the mean residence time, one needs to properly average the two durations above. The weights to perform the average are given by the probabilities $A_{\mathrm{ON}}$ and $A_{\mathrm{OFF}}$ that the system has just settled in these configurations. The residence time of a single TF being bound is then given by

$$T_{\mathrm{TF}} = \frac{e^{\delta}}{k_-} A_{\mathrm{ON}} + \frac{1}{k_-} A_{\mathrm{OFF}}. \tag{S15}$$

The probabilities $A_{\mathrm{ON}}$ and $A_{\mathrm{OFF}}$ are proportional to the product of i) the probability $W$ to find the system with a given binding site unoccupied, and ii) the rate of TF binding $k_+$. The probabilities $W$ for a given binding site being unoccupied can be calculated from the partition function $Z$ for $n$ binding sites (Eq. S7). After partitioning the states into Mediator `ON` and `OFF`, the resulting probabilities are proportional to the sum over all configurations of a $n-1$ system (Eq. S9):

$$\begin{aligned} W_{\mathrm{ON}} &= \frac{1}{Z} e^{L} \left(1 + c e^{\epsilon} e^{\delta}\right)^{n-1} \\ W_{\mathrm{OFF}} &= \frac{1}{Z} \left(1 + c e^{\epsilon}\right)^{n-1} \end{aligned} \tag{S16}$$

We can then express $A_{\mathrm{ON}}$ and $A_{\mathrm{OFF}}$ as

$$\begin{aligned} A_{\mathrm{ON}} &= k_{+} W_{\mathrm{ON}}/A \\ A_{\mathrm{OFF}} &= k_{+} W_{\mathrm{OFF}}/A, \end{aligned} \tag{S17}$$

where $A$ is normalization constant that ensures $A_{\mathrm{ON}} + A_{\mathrm{OFF}} = 1$. It follows that $A_{\mathrm{ON}} = W_{\mathrm{ON}}/(W_{\mathrm{ON}} + W_{\mathrm{OFF}})$ and $A_{\mathrm{OFF}} = W_{\mathrm{OFF}}/(W_{\mathrm{ON}} + W_{\mathrm{OFF}})$. Finally, by plugging these expressions into Eq. S15, we find that the residence time of a single TF on a binding site is given by

$$T_{\mathrm{TF}} = \frac{1}{k_{-}} \frac{W_{\mathrm{ON}} e^{\delta} + W_{\mathrm{OFF}}}{W_{\mathrm{ON}} + W_{\mathrm{OFF}}}. \tag{S18}$$

Numerical results show that our assumption about time-scale separation in this model is a valid approximation (Fig 2A, 2C).

Using the expression for the residence time above, we derive the lowest achievable $T_{\mathrm{TF}}$ in different scenarios. Since $T_{\mathrm{TF}}$ increases with the stability of complexes (i.e., by increasing $\alpha = e^{\delta}$), we can calculate the lower bound on $T_{\mathrm{TF}}$ by using the smallest possible $\alpha$, namely $\alpha = 1$ at fixed concentration and $\alpha = \alpha_{\mathrm{min}}$ (Eq. S13) at fixed expression $E_0$. We thus obtain the following $\min T_{\mathrm{TF}}$:

$$\min T_{\mathrm{TF}} = \frac{1}{k_{-}^{S}}, \qquad \text{for fixed } c, \tag{S19}$$

$$\min T_{\mathrm{TF}} = \min T_{\mathrm{TF}} = \frac{1}{k_{-}^{\mathrm{S}}} \cdot \frac{1}{1 + E_0(\alpha_{\mathrm{min}}^{-1} - 1)}, \qquad \text{for fixed } E. \tag{S20}$$

**Hill-type models.** As the number of possible equilibrium models for enhancer regulation is unlimited, let us consider at least one alternative to MWC models: Hill-type models. In these models, the presence of Mediator is not required to mediate the stabilization of TFs through links. Only the description of the TF interactions with DNA and the TF interaction with each other is necessary. In that scenario, linking could occur between neighbouring bound TFs (1D chain), between any pair of bound TFs, or via some other intermediate interaction scenario. As in the MWC model we considered, the creation of a link would lead to a multiplicative decrease in unbinding rate of TF by a factor of $\alpha$.

Since in Hill-type models we no longer have a two-state Mediator that naturally dictates what "active" enhancer (and thus expression) means, we need to revise our definition of expression. There are multiple possible definitions specifying the TF binding / linking configurations that lead to productive expression, i.e., are considered as effective `ON` states. The only constraint in order to preserve the proof-reading mechanisms and the high specificity advantage is that expression has to occur from states in which TFs are not only bound but also linked. For example, it could be i) all states that have at least one link, ii) only the state where all possible links are established, or iii) some other similar combination.

Let us show an example of non-equilibrium extension of a Hill-type model. We consider a 1D chain model where links can be established only between neighbouring bound TFs. In the non-equilibrium version of this model, links are not immediately created but are established with finite rate $k_{\text{link}}$. As in our MWC model extension, in the limit of $k_{\text{link}} \to \infty$, the states that differ only in the presence or absence of a link collapse into a single state. This occurs because as $k_{\text{link}}$ increases, the transitions from unlinked to linked state become much faster, until the two states are indistinguishable. Fig S.II shows an example for $n = 2$ binding sites. In the equilibrium limit at large $k_{\text{link}}$, assuming expression occurs only from the linked TF state, it is straightforward to write down the partition function, show that it predicts a Hill function with $n = 2$ for the induction curve, and that parameter $\alpha$ is directly related to the cooperative energy of interaction carried by the "link".

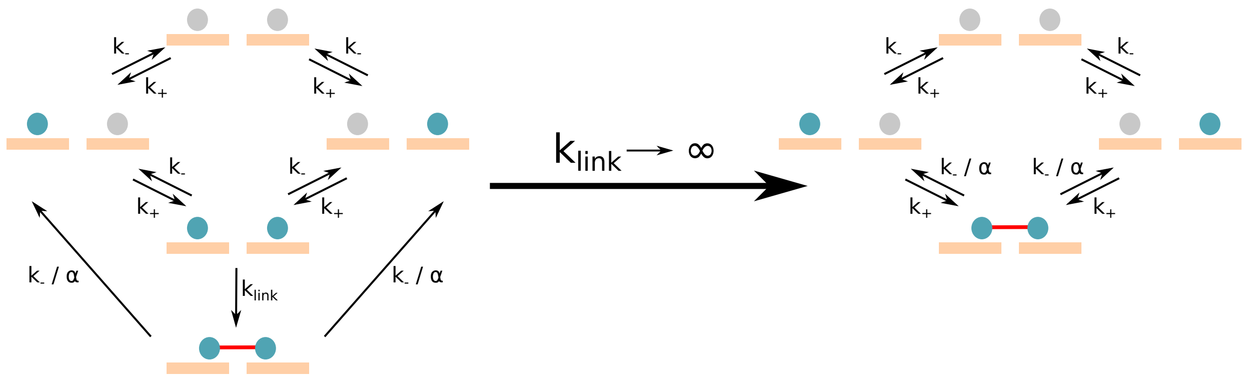

Fig S.II: **Non-equilibrium model extension of a Hill-type model.** A comparison between schemes of non-equilibrium extension (left) and equilibrium (right) Hill-type model for $n = 2$. When $k_{\text{link}} \to \infty$, the two models collapse. In the example of this model, links can be established only between two neighbouring bound TFs (red line).

#### 1.4 Regulatory phenotypes

**Expression** $E$ is the normalized expression level of a gene expressed under the control of the modeled enhancer. We compute expression as the fraction of time the Mediator is in `ON` state, that is $E = \langle s_M \rangle$. The average is taken over the stationary distribution of the master equation (except where we study transient effects, as in main text Fig 4). In practice, this means that we first compute the stationary solution $\mathbf{V}$ of Eq. S1: $\mathrm{d}\mathbf{V}/\mathrm{d}t = \hat{M}\mathbf{V} = 0$. We then marginalize $\mathbf{V}$ to obtain $E = \sum_{j=1}^{m} V_j I(s_M = 1)$, where the sum is taken over all the states of the Markov chain and $I(s_M = 1)$ is the indicator function which is 1 if the Mediator is `ON` in state indexed with $j$ and zero otherwise. As in the equilibrium limit, the expression is bounded: $E \in (\frac{k_+}{\kappa_-}, 1)$.

We expect that functional enhancers lead to high expression when TF concentration is high, which, in our model, should correlate with high occupancy of TFs on the specific BSs in the enhancer. We thus require the Mediator to be `ON` with high probability (typically $E \sim 0.5$, although we also consider in the main paper scenarios where $E$ can be smaller).

**Specificity** $S$ is the ratio between the level of expression from a functional enhancer (i.e., enhancer that contains $n$ specific BSs), and expression from a random piece of sequence, $S = E^{\mathrm{S}}/E^{\mathrm{NS}}$. High specificity of regulation ($S > 1$) is generally realized as a collective state of many bound TFs interacting with the Mediator. As in the equilibrium limit, specificity is bounded $S \in (1, \kappa_-/\kappa_+)$. In addition, when the specific expression is fixed $E \neq 1$ (as in Fig. 2C), the upper bound is adjusted by a factor of $E$, such that $\max S_{\text{fixed E}} = E\kappa_-/\kappa_+$. In our model, we estimate specificity by independently computing the expression $E$ for specific and non-specific site (i.e., for two different values for the unbinding rate, $k_-^{\mathrm{S}}$ and $k_-^{\mathrm{N}S}$), then taking their ratio.

We expect specificity to be as high as possible. Indeed, high specificity allows for accurate binding and control: first, it ensures that most of the TFs are not sequestered stably on random sequences; second, this further ensures that non-specific binding of TFs to non-cognate regulatory regions in the genome does not lead to erroneous gene expression, also known as transcriptional crosstalk. Given the high relative excess of possible non-specific binding configurations in the genome that outnumber binding configuration in the cognate regulatory region by thousands or millions, specificity should numerically be as high as possible.

**Residence time** $T_{\rm TF}$ is the average time that a TF spends bound to its specific BS. As in the equilibrium limit, the shortest TF residence time is obtained in absence of any stabilizing links, i.e.,

$$\min T_{\rm TF} = \frac{1}{k_-^{\rm S}}. \tag{S21}$$

In other words, the minimal TF residence time is determined by the unbinding rate of an isolated TF. Furthermore, when we consider the enhancer at a fixed specific expression $E^{\rm S} = E_0$ (as in Fig. 2C), this value gets adjusted to

$$\min T_{\rm TF} = \frac{1}{k_-^{\rm S}} \cdot \frac{1}{1 + E_0(\alpha_{\rm min}^{-1} - 1)}, \tag{S22}$$

with $\alpha_{\rm min} = \left(\frac{\kappa_-}{\kappa_+}\frac{E_0}{1-E_0}\right)^{1/n}$. This bound is obtained from the EQ model given the constraint for $E^{\rm S} = E_0$ (see Chapter 1.3). In our model, we computed the mean TF residence time directly from transition matrix $\hat{M}$ of the system (Eq. S1). More details about the residence time distributions and the moments can be found in Section 1.2.

Overall, we expect TF residence time to be small. Indeed, small residence time should provide better responsiveness to regulatory elements and lower the noise in gene expression. Furthermore, small residence time is consistent with recent single-molecule measurements. Since residence time is expected to increase with increasing stability of complexes (i.e., by increasing $\alpha$ and $k_{\rm link}$), there should be some trade off residence time and specificity.

**Sensitivity** $H$ refers to the effective steepness of the steady-state input/output curve that maps out the gene expression level as a function of the TF concentration.

We compute sensitivity $H$ as the slope of the induction curve (expression $E$ vs concentration of TFs $c$ on functional enhancers containing specific BSs with off-rate $k_-^{\rm S}$) at half maximum expression $E$, i.e., $H = 4\frac{c_{1/2}}{E_{\rm max}}\frac{{\rm d}E}{{\rm d}c}|_{c=c_{1/2}}$, where $\frac{{\rm d}E}{{\rm d}c}|_{c=c_{1/2}}$ represents the derivative of expression with respect to the concentration, taken at concentration $c_{1/2}$ where expression reaches half its maximum value $E_{\max}$. The normalization factor $4\frac{c_{1/2}}{E_{\max}}$ ensures that $H$ is properly bounded between 1 and $n$. Indeed, for Hill-like functions, $E(c) = c^h/(c^h + K^h)$, the defined sensitivity corresponds exactly to the Hill coefficient, $H = h$.

We construe sensitivity $H$ broadly, in terms of its functional effect on the shape of the induction curve regardless of the underlying molecular mechanism. Mechanisms giving rise to high sensitivity could be very diverse, for example: additional energy contribution due to a physical interaction of two TFs at the binding site, as in thermodynamic models of regulation; or a collective effect of competition of TF binding with nucleosomes, as in the MWC-like model proposed by Mirny et al. *PNAS* **107** (2010); or as a result of positive auto-regulation of a transcribed gene; or as a result of kinetic regulatory models out-of-equilibrium; or any other alternative that can increase the steepness of the induction curve beyond $H = 1$.

**Mean protein number $P$** represents the amount of protein, assuming protein dynamics is a deterministic consequence of the enhancer state. Its dynamics are governed by:

$$\frac{\mathrm{d}P}{\mathrm{d}t} = k_P R(t) - \frac{P}{T_P}, \tag{S23}$$

where $P$ represents the protein number, $k_P$ and $1/T_P$ the protein production and degradation rate, respectively, and $R(t)$ the enhancer state: 1 for `ON` and 0 for `OFF`.

$R(t)$ is a random variable whose stochastic realizations can be computed using stochastic simulation. To this end, we evolve the system state using a propagator, i.e., the formal solution of Eq. S1, which gives the conditional probability that the system will be found in some state after time $\Delta t$, given its current state. The enhancer state $R(t)$ is updated after each $\Delta t$ by randomly drawing a binary random number according to the probabilities computed using the propagator. Mathematically, the vector of probabilities $\mathbf{W}$ to go from state $j$ to any other state after time $\Delta t$ can be written as:

$$\mathbf{W}(\Delta t) = \exp(\hat{M}\Delta t)\ \mathbf{I}_j, \tag{S24}$$

where $\hat{M}$ is the transition matrix (see Eq. S1), $\mathbf{I}_j$ a vector of zeros with value 1 at $j$-th entry, representing the $j$-th state, and "exp" represents matrix exponential (see the formalism in Section 1.5 for details). We compute $R(t)$ at fixed $\Delta t$ time steps, with $\Delta t \ll T_{\mathrm{M}}$ and $\Delta t \ll T_{\mathrm{TF}}$, making sure that no representative `ON` state is missed.
After generating a stochastic realization $R(t)$, we solve the Eq. S23 using standard ODE solvers using $k_p = 1$ and $T_P = 3.6 \cdot 10^6$. Assuming $1/k_-^S = 1$ s, then $T_P = 10$ h. Results in Fig. 4C (middle- and bottom panel) represent the mean and standard deviation over 1000 replicates for different stochastic realizations of the enhancer state.

**Noise in protein number** $N$ represents the variability in protein expression levels due to random enhancer state switching. $N$ is defined as $\sigma_P/P$, where $P$ and $\sigma_P$ represent the mean and the standard deviation of protein number, respectively. For calculation of dynamical trace of the noise in Fig 4C, we followed the same procedure as for mean protein number (above).

##### 1.5 Noise propagation

**Telegraph model** Here, we briefly review some general results regarding the telegraph model or 2-state model that includes protein production and degradation with constant rates $k_P$ and $\gamma_P = 1/T_P$. The temporal evolution of the central moments can be derived from the master equation. The mean protein number $P$ and the mean gene activity $E$ satisfy the following equations

$$\begin{cases} \dfrac{d}{dt}P(t) = k_P E(t) - \gamma_P P(t) \\ \dfrac{d}{dt}E(t) = -(\kappa_+ + \kappa_-)E(t) + \kappa_+. \end{cases} \tag{S25}$$

At steady state ($\frac{d}{dt}P = 0$ and $\frac{d}{dt}E = 0$), the mean protein number and the mean activity is simply given by $P = P_0 E$ with $P_0 = k_P/\gamma_P$ and $E = \kappa_+/(\kappa_+ + \kappa_-)$. Similarly, the covariances satisfy the following set of equations

$$\begin{cases} \dfrac{d}{dt}\sigma_P^2(t) = -2\gamma_P\sigma_P^2(t) + 2k_P\sigma_{PE}(t) + \gamma_P P(t) + k_P E(t) \\ \dfrac{d}{dt}\sigma_{PE}(t) = -(\gamma_P + \kappa_+ + \kappa_-)\sigma_{PE}(t) + k_P\sigma_E^2(t), \end{cases} \tag{S26}$$

where the gene state variance $\sigma_E^2(t)$ is directly determined from the evolution of the mean $E(t)$, i.e. $\sigma_E^2(t) = E(t)(1 - E(t))$. This follows immediately from the fact that one state being occupied (either the active or inactive one) necessarily implies that the other is empty. Thus $\sigma_E^2$ must be the binomial variance at all time. Solving the equations S26 at steady state leads to

$$\sigma_P^2 = P_0 E + P_0 \sigma_{PE}$$
$$\sigma_{PE} = P_0 \frac{\gamma_P}{\gamma_P + \kappa_+ + \kappa_-} \sigma_E^2.$$

It follows that the protein variance is given by

$$\sigma_P^2 = P_0 E + P_0^2 E(1 - E)\Phi(T_P/T_E) \tag{S27}$$

where $\Phi(x) = 1/(1 + x) \in [0, 1]$ is a noise averaging/filtering function that determines the amount of propagated switching noise at the level of proteins by comparing the two relevant time scales of the system, namely the mean protein life time $T_P = 1/\gamma_P$ and the switching correlation time $T_E = 1/(\kappa_+ + \kappa_-)$. The first term $P_0 E$ in Eq. S27 corresponds to the Poisson variance resulting from the birth and death of proteins, while the second term stems from the propagation of the switching binomial variance

$$\left(\frac{dP}{dE}\right)^2 \sigma_E^2 \cdot \Phi(T_P/T_E) = P_0^2 \underbrace{E(1 - E)}_{\text{binomial variance}} \Phi(T_p/T_e). \tag{S28}$$

In the limit of fast and slow gene switching respectively, the noise filtering function reduces to

$$T_P \gg T_E \quad \lim_{x\to\infty} \Phi(x) = 0$$
$$T_P \ll T_E \quad \lim_{x\to 0} \Phi(x) = 1.$$

As we will see later, Eq. S27 remains valid for all the considered enhancer models, although the functional form of $\Phi$ will now depends on the details of the kinetic scheme considered.

The protein noise at steady state is thus generally given by

$$N^2 = \frac{\sigma_P^2}{P^2} = \frac{1}{P} + \frac{1-E}{E}\Phi(T_P/T_E) \simeq \frac{1-E}{E}\frac{T_E}{T_E+T_P}, \tag{S29}$$

where in the last equality we use the filtering function of the 2-state model and we drop the Poisson noise term $1/P$ assuming large number of proteins. It turns out that the last expression still provides an excellent approximation for the amount of propagated noise in the case of sophisticated $n$-state model, provided we use a good proxy for the switching correlation time $T_E$, which we will address below.

**General $m$-state enhancer model** For any $m$-state model of gene activity[1] where protein production and degradation occur as Poisson processes with constant rates $k_P$ and $\gamma_P$, the protein noise will satisfy the same functional form as the 2-state model (Eq. S29). In this general context, the gene mean activity $E$ is defined as the total mean occupancies of all the gene states $i$ allowing protein production, namely $E = \sum_{i=1}^{m_p} v_i$, with $1 \leqslant m_p < m$ the number of producing states and $v_i$ the mean occupancy of state $i$. It turns out that the switching noise can still be obtained by propagation of the binomial variance $\sigma_E^2 = E(1-E)$ multiplied by some noise filtering function $\Phi_m \in [0,1]$ (Eq. S28). The only difference for a $m$-state model of gene activity lies in the noise filtering function $\Phi_m$ that depends on the kinetic rates and topology of the gene state transition network, i.e. the $m \times m$ state transition matrix $M$ of the model. Starting from the equations for the first and second moment derived from the master equation (Eq. S1), we generalize the noise filtering function obtained for the 2-state model (Eq. S27) to an arbitrary number of gene states and transitions. The first moment equations are given by

$$\begin{cases} \dfrac{d}{dt}P(t) = k_P \mathbf{v_p^t} \mathbf{V}(t) - \gamma_P P(t) \\ \dfrac{d}{dt}\mathbf{V}(t) = \hat{M}_r \mathbf{V}(t) + \mathbf{f}, \end{cases} \tag{S30}$$

where $\mathbf{V}(t) = (v_1(t), v_2(t), ..., v_{m-1}(t))^{\mathbf{t}}$ is the vector of mean occupancies, or equivalently the probability to find the system in each individual gene state $i \in \{1, .., m-1\}$. The

[1]gene states described by a continuous time Markov process with linear propensity functions

vector $\mathbf{v_p^t} = (1, ..., 1, 0, ..., 0)$ defines which gene states permit protein production, such that $E(t) = \mathbf{v_p^t V}(t)$. The operator $\hat{M}_r$ is obtained by reduction of the original transition matrix $\hat{M}$

$$\hat{M}_r = A_1 \hat{M} A_0,$$

where $A_1$ and $A_0$ are the following $(m-1) \times m$ and $m \times (m-1)$ matrices

$$A_1 = \begin{pmatrix} 1 & & & 0 \\ & \ddots & & \vdots \\ & & 1 & 0 \end{pmatrix} \quad A_0 = \begin{pmatrix} 1 & & \\ & \ddots & \\ & & 1 \\ -1 & \dots & -1 \end{pmatrix}.$$

The vector $\mathbf{f}$ is given by the first $m-1$ terms of the last column of $M$. The reduction above is necessary to later on invert the $\hat{M}_r$ operator. Indeed, the transition matrix $\hat{M}$ is degenerate by construction and has a single zero eigenvalue corresponding to the steady state solution (provided the system is ergodic), which follows from $\sum_i \hat{M}_{ij} = 0 \ \forall j$ (that ensures conservation of probability). The occupancy of the last state $m$ is thus given by $v_m(t) = 1 - \sum_{i=1}^{m-1} v_i(t)$. Assuming steady-state, the gene state occupancies $\mathbf{V}$ are calculated from

$$\hat{M}_r \mathbf{V} + \mathbf{f} = \mathbf{0} \tag{S31}$$

and the mean protein is given by $P = k_P \mathbf{v_p^t V} / \gamma_P = P_0 E$ as in the 2-state model. Similarly, the time evolution of the covariances can be derived from the master equation and are given by

$$\begin{cases} \dfrac{d}{dt}\sigma_P^2(t) = -2\gamma_P \sigma_P^2(t) + 2k_P \mathbf{v_p^t} \boldsymbol{\sigma}_{PV}(t) + \gamma_P P(t) + k_P \mathbf{v_p^t V}(t) \\ \dfrac{d}{dt}\boldsymbol{\sigma}_{PV}(t) = -(\gamma_P \hat{I} - \hat{M}_r)\boldsymbol{\sigma}_{PV}(t) + k_P \hat{S}(t)\mathbf{v_p}, \end{cases} \tag{S32}$$

where $\boldsymbol{\sigma}_{PV}(t) = \big(\sigma_{P1}(t), \sigma_{P2}(t), ..., \sigma_{P(m-1)}(t)\big)^{\mathbf{t}}$ is the vector of covariances between the protein and each gene state, $\hat{I}$ the identity matrix, $\hat{S}(t)$ the covariance matrix of the gene states. Here again, it is important to realize that each gene state can only be occupied if all the others are empty. Thus, the covariance matrix $\hat{S}(t)$ is the multinomial covariance for a single trial, which is fully determined by the mean occupancies $\mathbf{V}(t)\ \forall t$

$$\hat{S}_{ij}(t) = \begin{cases} v_i(t)(1 - v_j(t)) & \text{for } i = j \\ -\, v_i(t)v_j(t) & \text{for } i \neq j. \end{cases} \tag{S33}$$

Solving Eqs S32 at steady state, we find

$$\begin{aligned} \sigma_P^2 &= P_0 E + P_0 \mathbf{v_p^t} \boldsymbol{\sigma}_{PV} \\ \boldsymbol{\sigma}_{PV} &= P_0 (\hat{I} - \hat{M}_r/\gamma_P)^{-1} \hat{S} \mathbf{v_p}. \end{aligned} \tag{S34}$$

By rearranging the steady state solutions (Eq. S34), we recover an expression for the protein variance $\sigma_P^2$, which is similar to the one derived before for a simple switch (cf. Eq. S27):

$$\sigma_P^2 = P_0 E + P_0^2 E(1 - E)\Phi_m. \tag{S35}$$

The main difference is the filtering function $\Phi_m$ now given by

$$\Phi_m = \frac{1}{E(1-E)} \mathbf{v_p^t} (\hat{I} - T_P \hat{M}_r)^{-1} \hat{S} \mathbf{v_p}, \tag{S36}$$

with $T_P = 1/\gamma_P$ the mean protein lifetime as before. In Eq. S36, the binomial variance $\sigma_E^2 = E(1-E)$ can be written as follows

$$E(1{-}E) = \sum_{i=1}^{m_p} v_i \left(1 - \sum_{i=1}^{m_p} v_i\right) = \sum_{i=1}^{m_p} v_i(1{-}v_i) {-} 2 \sum_{i<j\leqslant m_p} v_i v_j = \sum_{i=1}^{m_p} \sigma_i^2 {+} 2 \sum_{i<j\leqslant m_p} \sigma_{ij} = \mathbf{v_p^t} \hat{S} \mathbf{v_p}.$$

Plugging the above expression for the binomial variance back in Eq. S36, we finally find the following expression for the filtering function

$$\Phi_m = \frac{\mathbf{v_p^t} \hat{F} \hat{S} \mathbf{v_p}}{\mathbf{v_p^t} \hat{S} \mathbf{v_p}} \quad \text{with} \quad \hat{F} = (\hat{I} - T_P \hat{M}_r)^{-1}. \tag{S37}$$

Of note, the $\hat{F}^{-1}$ operator is positive definite[2], which follows from the positive definiteness of $-\hat{M}_r$ and $T_P \geqslant 0$. In addition, the spectrum of $\hat{F}^{-1}$ is bounded from below, i.e. all its eigenvalues $\lambda_i \geqslant 1$. Thus, $\mathbf{v_p^t}\hat{F}\hat{S}\mathbf{v_p} \leqslant \mathbf{v_p^t}\hat{S}\mathbf{v_p}\ \forall T_P$ and the resulting filtering function $\Phi_m$ satisfies all the desired conditions, namely $\Phi_m \in [0,1]$ and

$$\lim_{T_P\to\infty} \Phi_m(T_P) = 0$$
$$\lim_{T_P\to 0} \Phi_m(T_P) = 1.$$

In addition, we recover the correct expression for the 2-state model, where $\hat{M}_r = -(\kappa_+ + \kappa_-) = -1/T_E$. Indeed, $\hat{F} = (1 + T_P/T_E)^{-1}$ and thus $\Phi_2 = T_E/(T_E + T_P)$ as expected.

**Propagated noise and effective correlation time** As we have shown above, all the $m$-state models lead to the same functional form for the mean and variance (Eq. S35), differing only in the noise filtering function $\Phi_m$. Although multiple time scales, given by the inverse spectrum of $-\hat{M}_r$, are involved in the noise filtering, we can define a single effective switching correlation time $T_E$ that preserves as well as possible the amount of propagated noise of the $n$-state model. We aim for a definition of $T_E$ that is independent of the value of $T_P$, such that the resulting $T_E$ characterizes the filtering for all $T_P$ well. One way of proceeding is to realize that in the case of the 2-state model, $T_E = T_P$ implies $\Phi = 1/2$. We can thus use this property to define $T_E$ such that $\Phi_m(T_P = T_E) = 1/2$. Based on Eq. S37, we can then solve the following equation to obtain an effective $T_E$

$$\frac{1}{2}\mathbf{v_p^t}\hat{S}\mathbf{v_p} = \mathbf{v_p^t}(\hat{I} - T_E\hat{M}_r)^{-1}\hat{S}\mathbf{v_p}. \tag{S38}$$

With such an effective $T_E$, the filtering function $\Phi_m$ is well approximated by

$$\Phi_m(T_P) \simeq \Phi(T_P/T_E) = \frac{T_E}{T_E + T_P}, \tag{S39}$$

[2]Although $\hat{M}_r$ or $\hat{F}$ are not necessarily symmetric, $-\mathbf{x^t}\hat{M}\mathbf{x} > 0\ \forall$ non-zero vector $\mathbf{x}$ and all the eigenvalues of $-\hat{M}_r$ are positive. These properties follow from the structure of the master equation transition matrix $\hat{M}$.

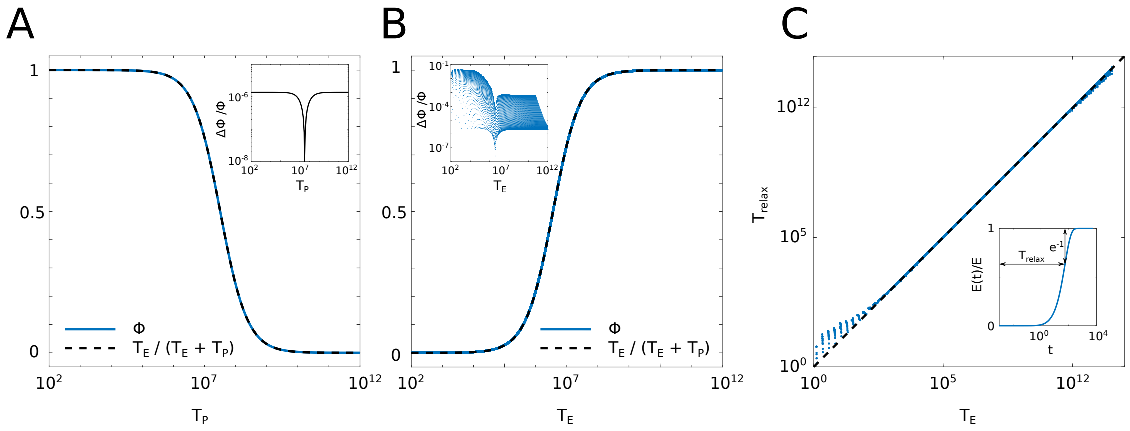

Fig S.III: **Effective correlation time determines propagated noise and relaxation time.** (A) Propagated noise fraction as a function of protein lifetime $T_P$ computed for parameters as model II in Fig 2A,C ($\alpha = 1.75 \cdot 10^4$, $k_{\text{link}} = 6.5 \cdot 10^{-2}$). The simple noise averaging function $T_E/(T_E + T_P)$, where $T_E$ is the effective correlation time, provides an excellent approximation to the true propagated noise $\Phi$. Indeed, the relative error (inset) remains small for the whole range of $T_P$. (B) Propagated noise fraction as a function of $T_E$ computed by probing the whole parameter space $\alpha \in (1, 10^8)$ and $k_{\text{link}} \in (10^{-5}, 10^5)$, at fixed $T_P$. The approximation $T_E/(T_E + T_P)$ captures the true propagated noise well over the full range of sampled models, as the relative error (inset) never exceeds 10%. (C) Relaxation time $T_{\text{relax}}$ as a function of effective correlation time $T_E$ for the whole parameter space as in (B). We estimated $T_{\text{relax}}$ from the temporal relaxation of the enhancer mean activity $E(t)$ to its steady state value $E$ (inset). To this end, we first solve Eq. S30 with $E(0) = 0$ to obtain $E(t)$ for each model. We then estimated $T_{\text{relax}}$ assuming $E(t)/E$ relaxes as $1 - \exp(-t/T_{\text{relax}})$, which is exact for the 2-state model. The resulting $T_{\text{relax}}$ matches $T_E$ well over the full range of sampled models. Thus our proposed effective correlation time $T_E$ is a good predictor of both propagated noise and mean relaxation time. In all panels, we used $n = 3$ binding sites for the models.

which is exact when $T_P = T_E$ and only slightly deviate from $\Phi_m(T_P)$ when $T_P > T_E$ or $T_P < T_E$, (Fig. S.IIIA,B).

Consequently, for all the enhancer models the propagated noise $N^2$ at the protein level is well approximated by

$$N^2 = \frac{1 - E}{E} \frac{T_E}{T_E + T_P}. \tag{S40}$$

In addition, the effective $T_E$ provides an excellent approximation for the mean relaxation time-scale of the models (Fig. S.IIIC).

##### 1.6 Effect of $\alpha$ and $k_{\text{link}}$ on regulatory phenotypes

To understand how varying $\alpha$ and $k_{\text{link}}$ affects regulatory phenotypes, we have a look at the phenotypes in the phase space $(\alpha, k_{\text{link}})$.

Fig S.IV shows the dependence of main regulatory phenotypes on $(\alpha, k_{\text{link}})$ for fixed concentration (left column) and fixed expression (right column). For fixed concentration, there exists only a narrow range that gives high specificity – around the point where specific expression is already large enough ($\sim 1$) while non-specific expression is still small ($\ll 1$). There, both residence time and sensitivity take relatively low values.

Meanwhile, for fixed expression, specificity increases with larger $\alpha$ and lower $k_{\text{link}}$. This is due to the fact that with increasing $\alpha$, the concentration required to reach fixed specific expression decreases, thus ensuring that non-specific expression stays low.

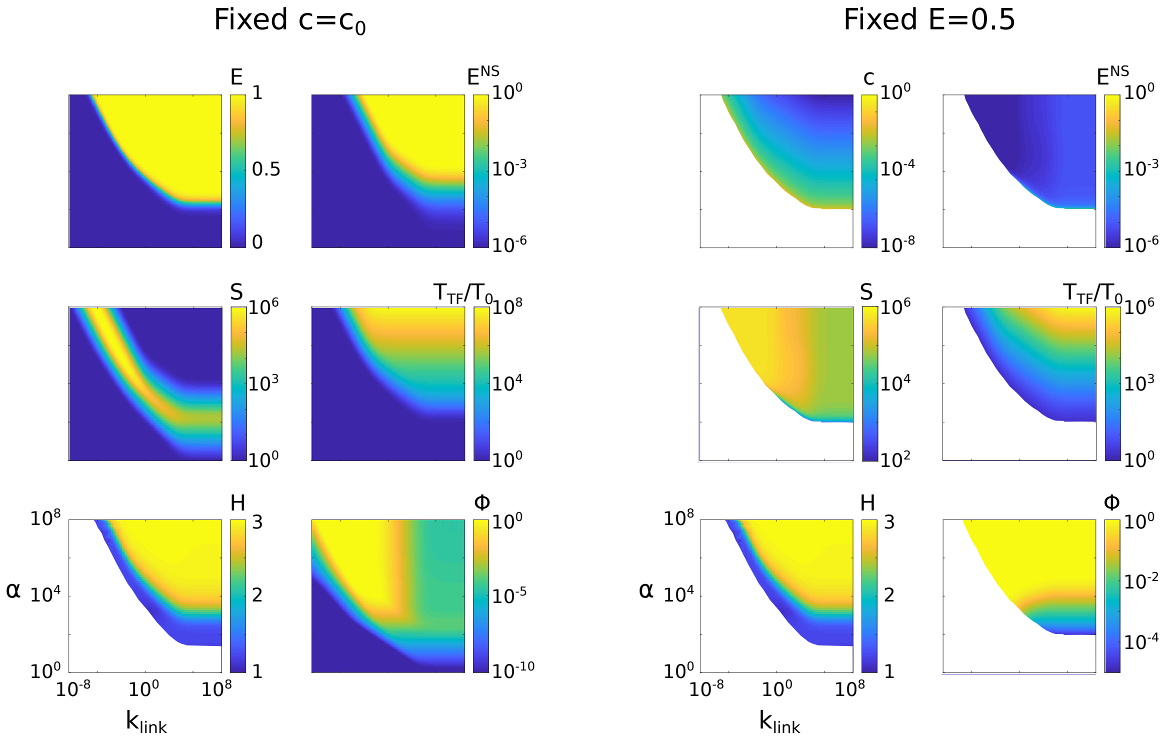

Fig S.IV: Expression from cognate enhancers containing $n = 3$ specific sites, $E^{\text{S}}$, and random DNA with nonspecific sites, $E^{\text{NS}}$, the TF residence time $T_{\text{TF}}$, specificity $S$, and sensitivity $H$ (in color) as a function of two parameters, $\alpha$ and $k_{\text{link}}$. Left and right column are showing regulatory phenotypes at fixed concentration $c = c_0$ and at fixed expression $E = 0.5$, respectively. For fixed expression, regions where $E = 0.5$ cannot be satisfied are colored white. Due to numerics, area of sensitivity $H$ where maximum expression (in the limit $c \to \infty$) is below $E < 10^{-3}$, is also colored white.

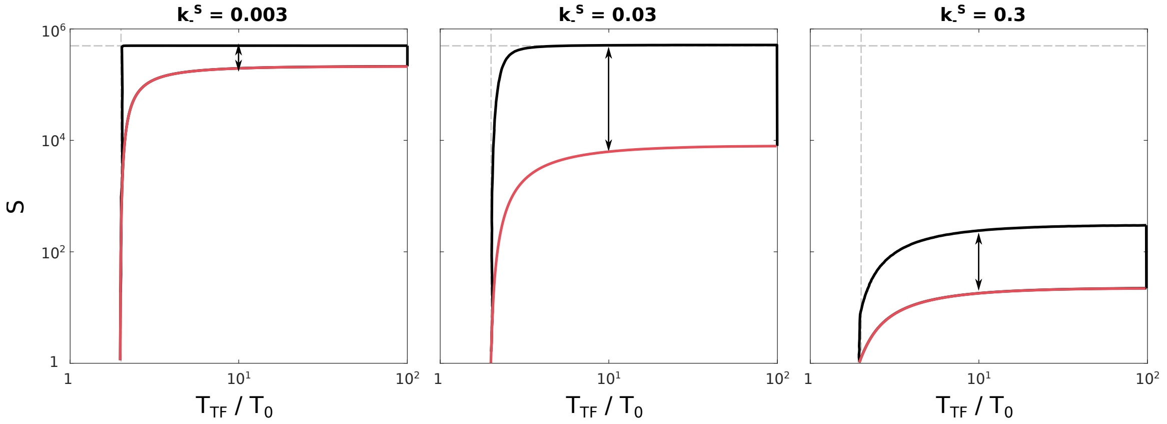

Fig S.V: Specificity as a function of TF residence time at fixed $E = 0.5$, showing how specificity gain (black arrow) changes for different values of $k_-^{\mathrm{S}}$ with fixed $k_-^{\mathrm{NS}} = 1$ at $n = 3$. EQ model solutions lie on the red line while the black/red envelope represents the space of solutions for NEQ model. The reference time is $T_0 = 1/k_-^{\mathrm{S}}$ which varies between the three figures. Dashed lines represent minimum residence time and maximum specificity.

#### 1.7 Effect of the unbinding rate ratio $k_-^{\mathrm{S}}/k_-^{\mathrm{NS}}$ on the specificity gain

In the main text we investigated how the maximum gain in specificity, $S_{\mathrm{NEQ}}/S_{\mathrm{EQ}}$, varies with the ratio of specific and non-specific unbinding rate, $k_-^{\mathrm{S}}$ and $k_-^{\mathrm{NS}}$, (Fig 3C). We identified an optimal value of $k_-^{\mathrm{S}}/k_-^{\mathrm{NS}}$ which maximizes this gain.

For smaller values of $k_-^{\mathrm{S}}/k_-^{\mathrm{NS}}$, NEQ model is at the bound of specificity, $S_{\mathrm{max}}$, with EQ model being close to it as well: see Fig S.V left. With increasing $k_-^{\mathrm{S}}/k_-^{\mathrm{NS}}$, the specificity of EQ model decreases, leading to an increase in the specificity gain (Fig S.V middle). The largest specificity gain is obtained when the maximum specificity in NEQ model is not bounded anymore but very close to it. With further increasing $k_-^{\mathrm{S}}/k_-^{\mathrm{NS}}$, the difference in specificity between NEQ and EQ model starts to decrease (Fig S.V right).

#### 2 SI Figures

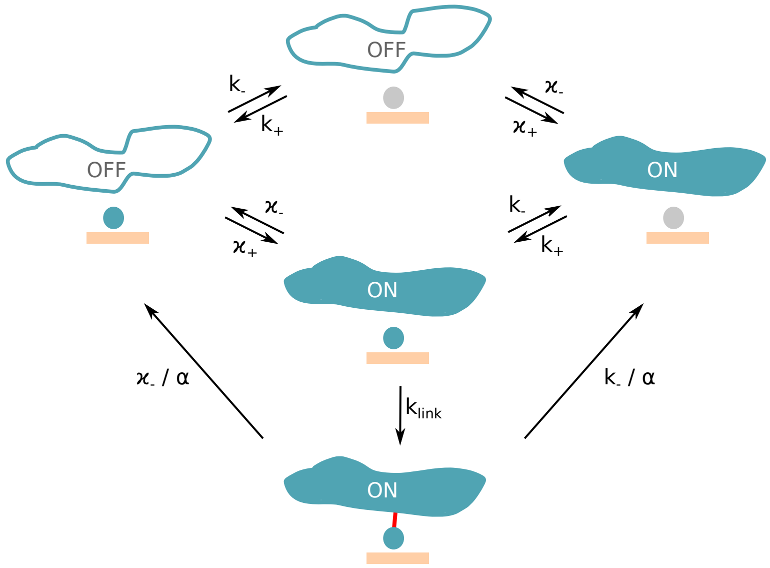

Fig S1: **Kinetic scheme of the non-equilibrium MWC-like model.** For simplicity, the scheme is illustrated for a single binding site, $n = 1$. TFs can bind to the specific binding site and Mediator can switch into `ON` state; when a TF is bound it can form a link only if a Mediator is found in `ON` state. The link decreases the unbinding rate of both linked TF and Mediator by a factor of $\alpha$. The link is removed either when the Mediator switches `OFF` or when the linked TF unbinds. With increasing number of binding sites $n$, the number of possible states exponentially increases.

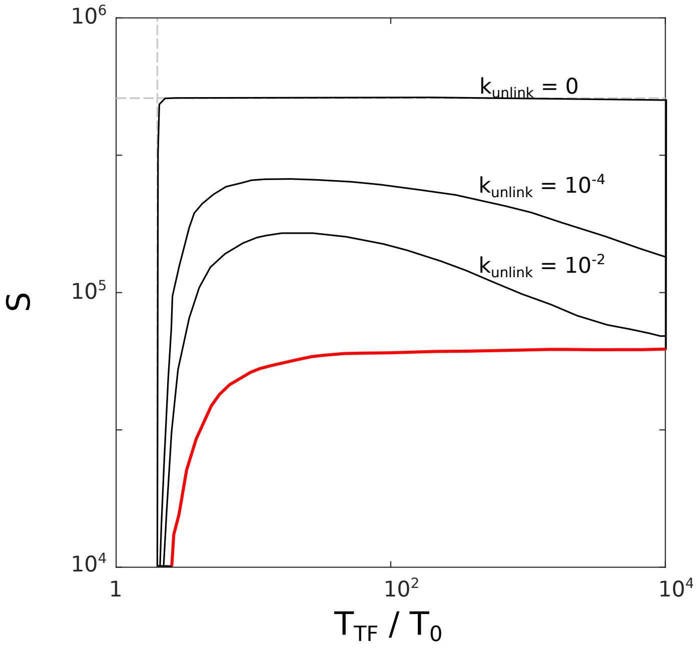

Fig S2: **Specificity effect of the nonzero unlinking rate.** The phase diagram of specificity, $S$, and mean TF residence time, $T_{\mathrm{TF}}$, for $n = 3$ binding sites and fixed $E = 0.5$, demonstrating the effect of unlinking rate. With increasing unlinking rate, the maximum specificity of the nonequilibrium model decreases. Furthermore, for non-zero unlinking rate, $k_{\mathrm{unlink}} > 0$, at larger TF residence times, the maximum specificity starts to decrease with TF residence time. This is qualitatively different than in case of a zero unlinking rate where maximum specificity never decreases with $T_{\mathrm{TF}}$. Each black envelope shows all solutions for varying $\alpha \in (1, 10^8)$ and $k_{\mathrm{link}} \in (10^{-5}, 10^{-8})$. Red curve represents equilibrium solutions at $k_{\mathrm{link}} \to \infty$, which do not vary with $k_{\mathrm{unlink}}$. We used $k_-^S = 0.01$ and $k_-^{NS} = 1$. See also Fig S9D.

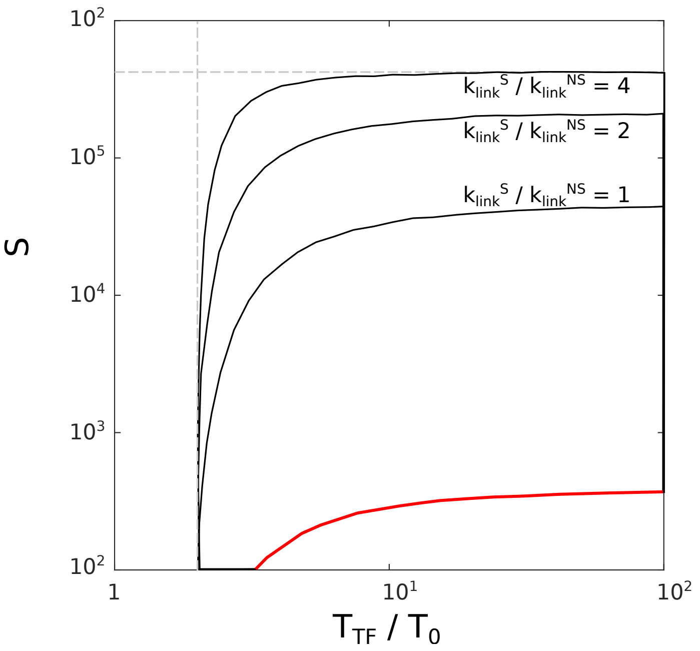

Fig S3: **Specificity gain due to sequence-specific linking rate.** The phase diagram of specificity, $S$, and mean TF residence time, $T_{\mathrm{TF}}$, for $n = 3$ binding sites and fixed $E = 0.5$, demonstrating the effect of different sequence-specific linking rates. We assume that the formation of link on a TF bound to a specific site is faster, by the indicated factor, $k_{\mathrm{link}}^{\mathrm{S}}/k_{\mathrm{link}}^{\mathrm{NS}}$, relative to the link formation when the TF is bound to a nonspecific site; this could happen, for instance, if the links are created by dedicated enzymes with their own DNA sequence binding preference. Large specificity increases are possible even at $k_{\mathrm{link}}^{\mathrm{S}}/k_{\mathrm{link}}^{\mathrm{NS}}$ not much larger than 1. We used $k_{-}^{\mathrm{S}} = 0.1$ and $k_{-}^{\mathrm{NS}} = 1$ (instead of the $k_{-}^{S} = 0.01$ used elsewhere). The ratio of $k_{\mathrm{link}}^{\mathrm{S}}/k_{\mathrm{link}}^{\mathrm{NS}} = 1$ represents the case in the main paper, without any linking sequence specificity. Each black envelope shows all solutions for varying $\alpha \in (1, 10^8)$ and $k_{\mathrm{link}} \in (10^{-5}, 10^{-8})$. Red curve represents equilibrium solutions, which do not vary with $k_{\mathrm{link}}^{\mathrm{S}}/k_{\mathrm{link}}^{\mathrm{NS}}$. Gray dashed lines show the analytically-derived bounds. The increase in $S$ due to linking specificity is seen only for ratios of $k_{-}^{\mathrm{S}}/k_{-}^{\mathrm{NS}}$ that are smaller that the optimal value (the ratio where $S_{\mathrm{NEQ}}/S_{\mathrm{EQ}}$ reaches a maximum, see Fig 3C). The reason is that the linking rate specificity affects only the nonequilibrium models, and for lower values of $k_{-}^{\mathrm{S}}/k_{-}^{\mathrm{NS}}$, the NEQ models already reach the maximum possible specificity, $\kappa_{-}/\kappa_{+}$; see Fig S.V. This means that for low values of $k_{-}^{\mathrm{S}}/k_{-}^{\mathrm{NS}}$, any additional linking specificity would not be able to increase NEQ enhancer specificity any further as it is already saturated. Linking specificity does not qualitatively change the overall conclusions, but can quantitatively boost specificity $S$.

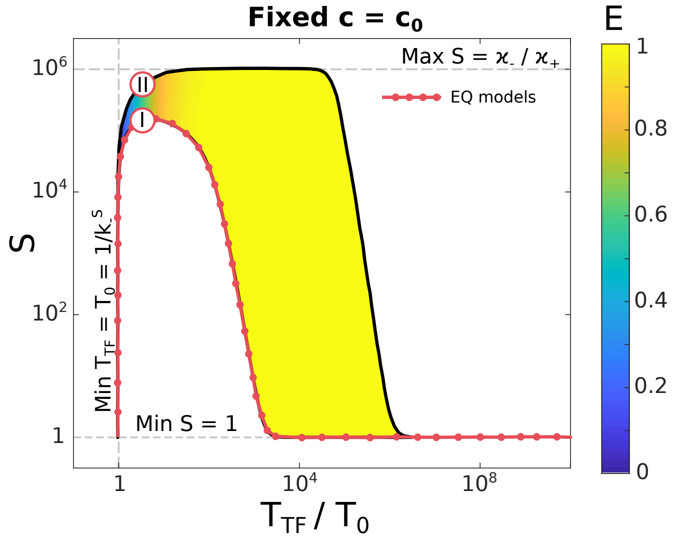

Fig S4: **Accessible space of regulatory phenotypes is similar for different number of binding sites.** Specificity, $S$, mean TF residence time, $T_{\text{TF}}$ (expressed in units in inverse off-rate for isolated TFs at their specific sites, $T_0 = 1/k_{-}^{\text{S}}$), and average expression, $E$ (color), for MWC-like models with $n = 5$ TF binding sites (main text Fig 2A showing $n = 3$), obtained by varying $\alpha$ and $k_{\text{link}}$ at fixed TF concentration, $c_0$. Equilibrium models fall onto the red line. As in the main text, two models with equal TF residence times, `I` (EQ) and `II` (NEQ), are marked for comparison. Dashed gray lines show analytically-derived bounds. The EQ model reaches higher specificity than for $n = 3$, making the space of solutions for $E < 1$ smaller. NEQ model solutions become limited by specificity ceiling, $S_{\max} = \kappa_{-}/\kappa_{+}$, and $S > 1$ solutions only exist for $T_{\text{TF}}/T_0 < 10^6$ (for $n = 3$ this was true for $T_{\text{TF}}/T_0 < 10^8$). Nevertheless, the accessible space of regulatory phenotype is qualitatively preserved for $n > 3$.

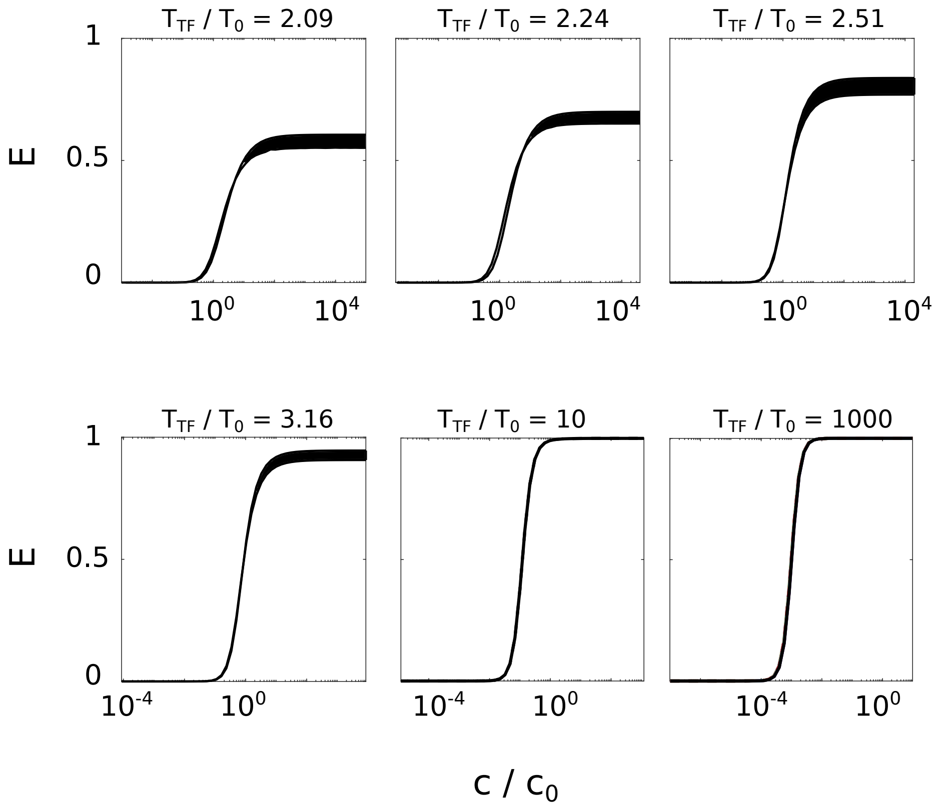

Fig S5: **Different models lead to indistinguishable induction curves from functional enhancers.** Induction curves for expression from functional enhancers (that contain specific binding sites) at fixed TF residence times (as indicated in the plot titles of different plots), for $n = 3$ and $E = 0.5$. In each plot with a given TF residence time, we find 20 different models with a range of specificities $S$, including the equilibrium model, and overlay their induction curves in black. Induction curves are nearly indistinguishable, with largest differences found for low residence times at large concentrations. The minimal achievable TF residence time is min $T_{\mathrm{TF}} = 1.98 T_0$.

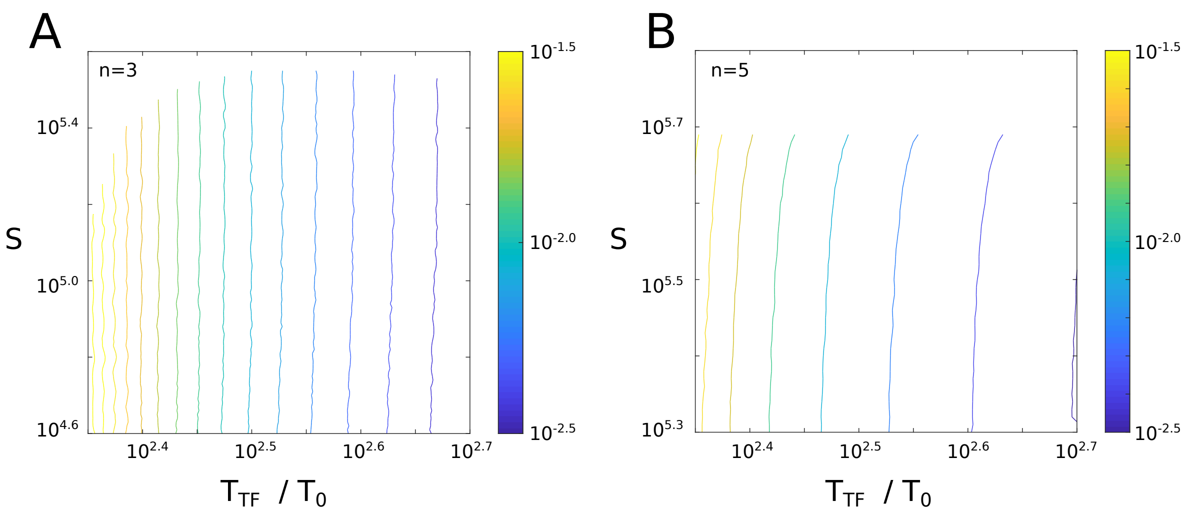

Fig S6: **Equi-concentration lines in enhancer phase diagrams at fixed expression are nearly vertical.** Lines of constant concentration (color) as a function of specificity and TF residence time for $n = 3$ (A) and $n = 5$ (B). Since all NEQ and EQ models have almost identical induction curves (Fig 2E and Fig S5), this implies that lines of constant concentration at fixed expression in the $S$ vs $T_{\mathrm{TF}}$ space (Fig 2C), are nearly vertical. This assumption holds very well for smaller values of $T_{\mathrm{TF}}$. With increasing residence time, lines of constant concentration start slightly tilting towards larger residence times. Additionally, for large specificity (close to the maximum specificity $\kappa_-/\kappa_+$) lines of constant concentration start to increase their curvature, especially at higher $n$.

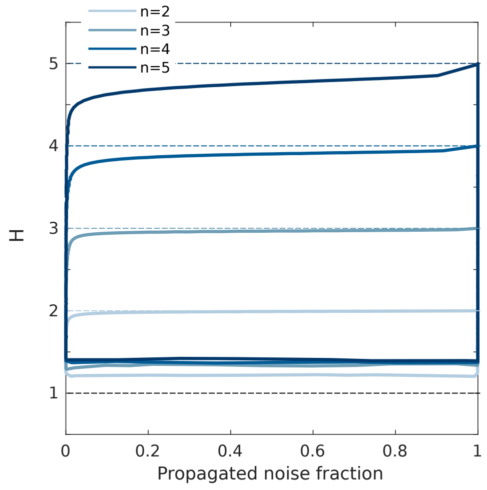

Fig S7: **Sensitivity and propagated noise fraction are uncorrelated.** Phase diagrams of sensitivity $H$ and propagated noise fraction, $T_E/(T_E+T_P)$ (see main text). Each envelope represents different value of $n$ (blue shade, legend); different models within each envelope are obtained by varying $\alpha \in (1, 10^8)$ and $k_{\mathrm{link}} \in (10^{-5}, 10^8)$, holding expression fixed at $E = 0.5$ by adjusting TF concentration. Almost all combinations of sensitivity and noise fraction are possible, indicating that these regulatory phenotypes are largely uncorrelated. The exception are highest possible sensitivities that are accessible only at higher values of propagated noise fraction.

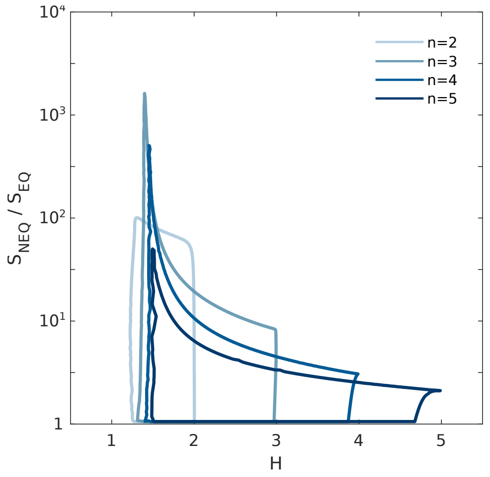

Fig S8: **Trade-off between optimal specificity and sensitivity.** Phase diagrams show the space of solutions of specificity gain, $S_{\text{NEQ}}/S_{\text{EQ}}$, and sensitivity, $H$. The envelopes were obtained by varying $\alpha \in (1, 10^8)$ and $k_{\text{link}} \in (10^{-5}, 10^8)$ at fixed expression $E = 0.5$ and various number of binding sites $n$ (blue shade, legend). Specificity gain is highest at lower residence times where the sensitivity is the lowest (Fig 3A). These results do not qualitatively vary with number of binding sites $n$, and show a trade-off between optimal specificity and high sensitivity gain.

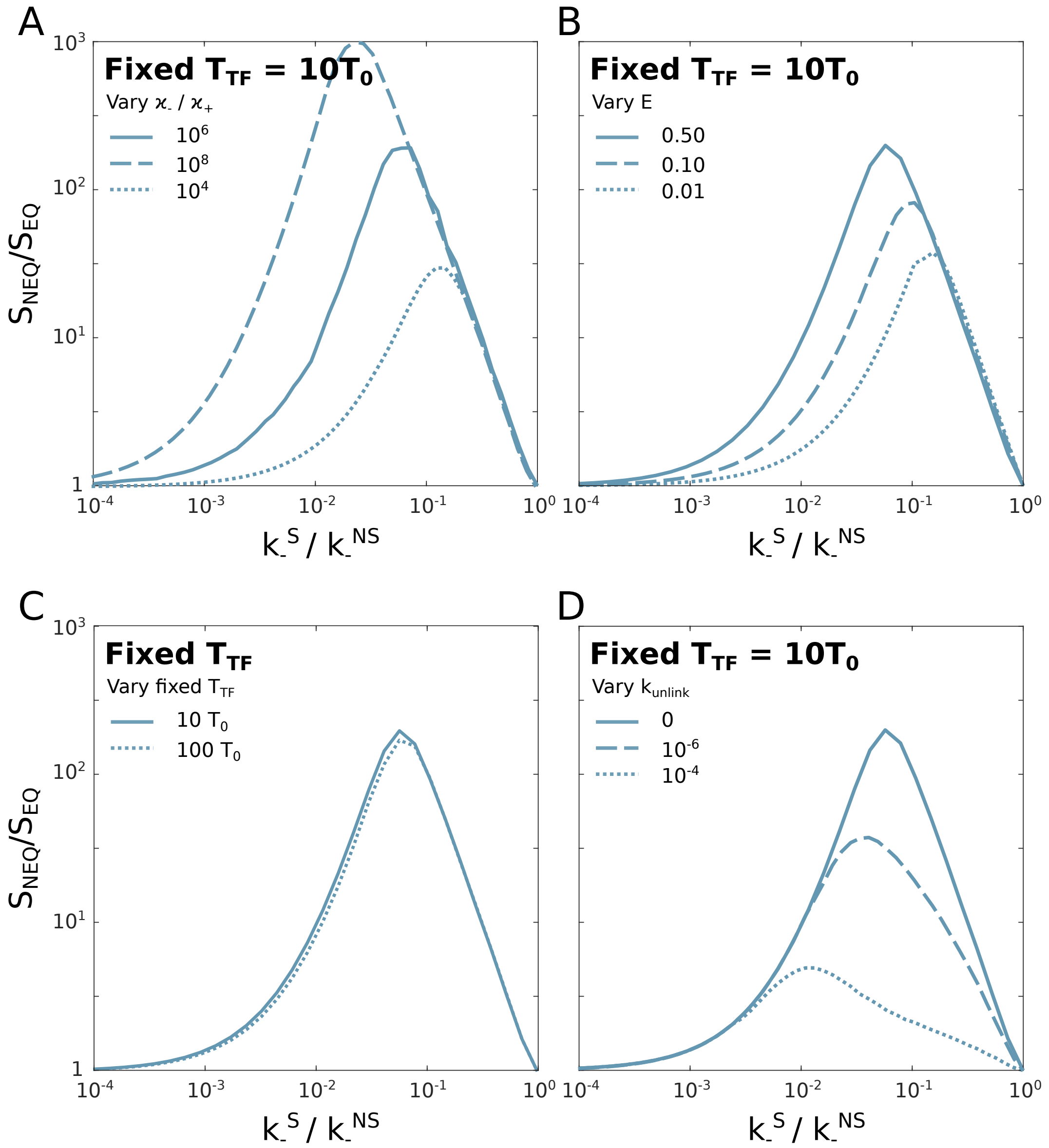

Fig S9: **Impact of kinetic and phenotypic parameters on optimal specificity gain.** Maximum gain in specificity as a function of a ratio of specific and non-specific unbinding rates, $k_-^{\mathrm{S}}/k_-^{\mathrm{NS}}$. Different plots show effects of different parameters that were varied: (**A**) ratio of Mediator switching rates $\kappa_-/\kappa_+$; (**B**) fixed expression $E$; (**C**) TF residence times at which the specificity is compared; and (**D**) unlinking rate $k_{\mathrm{unlink}}$. The strongest dependence is on Mediator switching rates, which set the upper bound for specificity, $S_{\mathrm{max}}$; when that increases, the maximum specificity gain also increases. Additionally, the value of fixed expression $E$ has a visible impact, similar as varying $\kappa_-/\kappa_+$. The residence time at which we compare specificity has a negligible role; for almost any value of the residence time, specificity gain does not vary much. This is due to the fact that for $T_{\mathrm{TF}} \gg T_0$, both EQ and maximum NEQ specificity do not significantly vary with $T_{\mathrm{TF}}$. Results show that for specificity gains right of the peak (i.e., larger $k_-^S/k_-^{NS}$), the ratio of Mediator switching rates $\kappa_-/\kappa_+$ does not play a visible role. The same is true for fixed expression $E$, and the TF residence time at which we compare specificity. When it is non-zero, $k_{\mathrm{unlink}}$ can strongly affect the specificity gain. For optimal gain, $k_{\mathrm{unlink}}$ must be much smaller than all other rates in the system.

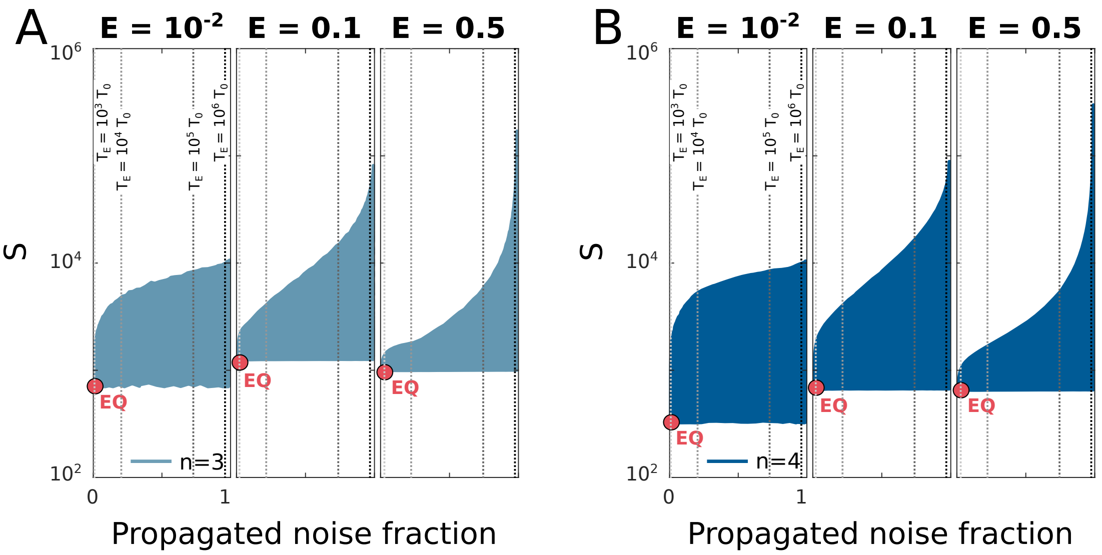

Fig S10: **Trade-off between optimal specificity and propagated noise.** Phase diagram of enhancer models for three different values of mean expression, $E$ (columns), shows specificity $S$ and fraction of variance in enhancer switching propagated to expression noise ($T_E/(T_E + T_P)$, see main text). Compact blue region for each $E$ shows all MWC-like models with $n = 3$ (**A**) and $n = 4$ (**B**) binding sites accessible by varying $\alpha \in (1, 10^8)$ and $k_{\text{link}} \in (10^{-5}, 10^5)$; equilibrium model ("EQ") with lowest noise is shown as a red dot. We have picked the $k_-^{\text{S}}/k_-^{\text{NS}}$ ratio to maximize the specificity gain $S_{\text{NEQ}}/S_{\text{EQ}}$: $k_-^{\text{S}}/k_-^{\text{NS}} \approx 0.06,\ 0.12$ for $n = 3,\ 4$, respectively (see Fig 3C). With these values, the specificity can increase but only at a cost of a large increase in correlation time $T_E$, implying high noise.
